# Automated Analysis of Low-Field Brain MRI in Cerebral Malaria

**DOI:** 10.1101/2020.12.23.424020

**Authors:** Danni Tu, Manu S. Goyal, Jordan D. Dworkin, Samuel Kampondeni, Lorenna Vidal, Eric Biondo-Savin, Sandeep Juvvadi, Prashant Raghavan, Jennifer Nicholas, Karen Chetcuti, Kelly Clark, Timothy Robert-Fitzgerald, Theodore D. Satterthwaite, Paul Yushkevich, Christos Davatzikos, Guray Erus, Nicholas J. Tustison, Douglas G. Postels, Terrie E. Taylor, Dylan S. Small, Russell T. Shinohara

**Affiliations:** Department of Biostatistics, Epidemiology, and Informatics, University of Pennsylvania; Mallinckrodt Institute of Radiology, Washington University in St. Louis; Department of Psychiatry, Columbia University Irving Medical Center; Blantyre Malaria Project, Kamuzu University of Health Sciences; Department of Radiology, Children’s Hospital of Philadelphia; Department of Radiology, Michigan State University; Tenet Diagnostics; Department of Diagnostic Radiology and Nuclear Medicine, University of Maryland School of Medicine; University Hospitals Cleveland Medical Center, Department of Radiology, Case Western Reserve University; Department of Paediatrics and Child Health, Kamuzu University of Health Sciences; Department of Psychiatry, University of Pennsylvania; Department of Radiology, University of Pennsylvania; Center for Biomedical Image Computing and Analysis (CBICA), Department of Radiology, University of Pennsylvania; Department of Radiology and Medical Imaging, University of Virginia; Division of Neurology, George Washington University/Children’s National Medical Center; College of Osteopathic Medicine, Michigan State University; Department of Statistics, University of Pennsylvania

**Keywords:** MRI, data integration, segmentation, Markov random field

## Abstract

A central challenge of medical imaging studies is to extract biomarkers that characterize disease pathology or outcomes. Modern automated approaches have found tremendous success in high-resolution, high-quality magnetic resonance images (MRI). These methods, however, may not translate to low resolution images acquired on MRI scanners with lower magnetic field strength. In low-resource settings where low-field scanners are more common and there is a shortage of radiologists to manually interpret MRI scans, it is critical to develop automated methods that can augment or replace manual interpretation, while accommodating reduced image quality. We present a fully automated framework for translating radiological diagnostic criteria into image-based biomarkers, inspired by a project in which children with cerebral malaria were imaged using low-field 0.35 Tesla MRI. We integrate multi-atlas label fusion, which leverages high-resolution images from another sample as prior spatial information, with parametric Gaussian hidden Markov models based on image intensities, to create a robust method for determining ventricular cerebrospinal fluid volume. We also propose normalized image intensity and texture measurements to determine the loss of gray-to-white matter tissue differentiation and sulcal effacement. These integrated biomarkers have excellent classification performance for determining severe brain swelling due to cerebral malaria.

## 1 Introduction

Malaria is a parasitic infection that results in more than 400,000 deaths annually. Although disease incidence has fallen steadily over the last decade, excess deaths due to malaria are expected to rise amid disruptions caused by the SARS-CoV-2 pandemic (World Health Organization, 2020). The great majority of malaria deaths occur in children living in sub-Saharan Africa (Luzolo and Ngoyi, 2019). A leading cause of mortality is cerebral malaria (CM), a serious complication of malaria infection clinically defined as coma in a person with malaria parasitemia (Idro et al., 2010). In children, CM has a case fatality rate of 15-20% despite optimal treatment (Dondorp et al., 2010). The pathophysiological mechanisms behind CM are not completely understood, though diffuse brain swelling, intracranial hypertension, and higher brain weight-for-age are commonly seen in fatal cases (Seydel et al., 2015). It is hypothesized that brain swelling, in conjunction with factors outside the central nervous system, plays a critical role in disease outcome. Brain magnetic resonance imaging (MRI), a non-invasive modality for assessing brain swelling, has been used to study CM pathogenesis (Looareesuwan et al., 2009), assess participants’ eligibility for enrollment in clinical trials (Kampondeni et al., 2018), and identify CM patients at highest risk of death. In the last two cases, the severity of swelling (cerebral edema) must be evaluated in real time, ideally by a trained on-call neuroradiologist.

MRI employs powerful electromagnetic fields to visualize brain structures and is a valuable tool for both disease diagnosis and prognosis. Recent trends in MRI hardware, including magnets with increasing field strengths of up to 7 Tesla (7T), have allowed capture of extremely detailed brain images. Concurrent with technology improvements, there has been an explosion of sophisticated and automated interpretation methods ranging from lesion segmentation in multiple sclerosis (Valcarcel et al., 2018; Valverde et al., 2017) or brain tumor (Gordillo et al., 2013; Mohsen et al., 2018) to the prediction of clinical outcomes in patients with psychosis (Nieuwenhuis et al., 2017). However, access to advanced MRI hardware and software-based interpretation methods is not consistent across the globe (Marques et al., 2019). In low-resource settings, challenges related to cost, infrastructure, and electric power supply may limit the availability of high field strength MRI (Ogbole et al., 2018; Latourette et al., 2010), sparking interest in lower field strength (< 0.5T) alternatives. Compared to high-field MRI, low-field MRI is more affordable and more easily sustained, but produces images with lower spatial resolution, since signal-to-noise ratio scales with magnet strength (Sarracanie and Salameh, 2020). Despite these challenges, low-field neuroimaging offers diagnostic value in situations where only moderate resolution is required for a clinical assessment, such as stroke (Bhat et al., 2020), infant hydrocephalus (Obungoloch et al., 2018), and CM (Kampondeni et al., 2013).

Automated interpretation methods are also relevant in resource-constrained settings as radiologists who can manually interpret brain scans are rare (Mollura et al., 2020), even when MRI is available. Because visual brain MRI interpretations are often unstructured and subjective, there is a growing demand for quantitative radiographic methods for CM (Potchen et al., 2013). An open question is whether modern image analysis pipelines—which were developed on higher resolution images—can be translated to low-resolution and lower quality scans, providing information from images acquired in low-resource settings.

We address this question by experimenting with a common pre-processing task in brain MRI analysis known as brain segmentation. Three-dimensional pixels (voxels) of the brain are identified and isolated from other voxels in the image. We show that popular surface-based methods such as the Brain Extraction Tool (Smith, 2002) and Brain Surface Extractor (Sandor and Leahy, 1997) perform well on 3T images, but poorly on the low-resolution images in our sample, despite parameter tuning.

In response, we present a novel, integrative framework that adapts image processing pipelines originally designed for high field strength scanners to images from low field strength scanners. In section 2, we introduce our neuroimaging dataset and describe the radiographic criteria currently used to assess severely increased brain volume (BV) in children with CM. In section 3, we describe a multi-atlas strategy using existing high-quality and publicly available brain imaging data to identify cerebral tissue in low field MRI. We then employ a Gaussian hidden Markov random field model to identify tissues within the brain based solely on the observed image intensities and spatial information. Finally, we adapt currently available image processing techniques to extract volume-, intensity-, and curvature-based biomarkers of severe BV, informed by imaging criteria used by expert raters. In section 4, we apply these biomarkers to obtain a predictive model of BV score in children with CM, validating model performance with an external dataset and nested cross-validation. In section 5, we discuss our findings and general principles for the analysis of images with limited spatial resolution. To our knowledge, this is the first fully automated method to assess biomarkers of severely increased BV in low field MRI.

## 2 Magnetic Resonance Imaging

Participants in this retrospective study were children (aged 6 months to 14 years old) admitted to the Blantyre Malaria Project, a long-term study of CM pathogenesis located in Blantyre, Malawi. All children had a Blantyre Coma Score of 2 or less, malaria parasitemia on peripheral blood smear, and no other known cause of coma. After clinical stabilization and beginning of intravenous antimalarial medications, participants were imaged with a General Electric 0.35T Signa Ovation MRI system (General Electric Healthcare, Chicago, Illinois). We considered two pulse sequences to highlight diverse tissue structures in the brain: a typical T1-weighted image exhibits brightest signal for fat, brighter signal for white matter than gray matter, and darkest signal for cerebrospinal fluid (CSF); in a T2-weighted image, CSF and fat are both bright and white matter is relatively dark. During the enrollment period, 100 participants were imaged. Five children were excluded from this analysis as they did not have both T1 and T2 sequences.

MRI acquisition parameters were not uniform across participants, nor were they uniform across modalities within a participant. For instance, while most participants had T1 and T2 scans that had high in-plane resolution in the axial dimension, many had a mixture of T1 and T2 scans that had high in-plane resolution in axial, coronal, and/or sagittal planes. A further challenge was that, in almost all images, the top of the brain was outside of the field of view; this was due to time constraints with respect to image acquisition. Finally, some images contained banding and other artifacts due to participant motion and technical factors.

The images were randomly partitioned into training (*n* = 48) and testing (*n* = 46) sets at the outset. In the training set, 3 radiologists scored every image; in the testing set, the same 3 radiologists and 5 additional radiologists scored every image. All radiologists were previously trained in the assessment of brain volume in pediatric CM, and scored images independently. We performed exploratory analyses and biomarker identification on the training data; a testing set of MRI scans was reserved to assess prediction performance.

### 2.1 Brain Volume Severity Score

MRI images were assigned a brain volume score ranging from 1 to 8 according to the degree of sulcal and cisternal effacement, grey-white matter delineation, and the presence of uncal herniation (Table 1) (Kampondeni et al., 2020; Potchen et al., 2012, 2018; Seydel et al., 2015). Patients with scores of 7 or greater are classified as having “highly increased” brain volume (BV), and are at higher risk of death compared to those with lower scores. Occasionally, radiologists assigned a score of 6.5, described as “moderate to severe,” when the images had features for both scores 6 and 7. Otherwise, they assigned scores that were integer-valued. BV scores were assigned based on all MRI images, including the T1 and T2 sequences as well as occasionally acquired diffusion weighted images. The automated interpretation methods developed here used only T1 and T2 sequences, as these were consistently available across all participants. As each participant’s BV score was assigned by multiple radiologists, the overall BV score for that participant was calculated as the median of these ratings.

**Table 1:** Radiological criteria used to assign brain volume (BV) scores and identify patients with severe brain swelling. A BV score ≥ 7 indicates severe or “highly increased” brain swelling. Radiologists occasionally assigned participants a score of 6.5 when the images had features for both BV = 6 and BV = 7; otherwise, they assigned integer-valued scores. The overall BV score for a given participant was the median BV score from all radiologists.

| Score | Definition |
| --- | --- |
| 1 | Severe atrophy markedly abnormal for age with diffuse prominence of the cisternae and sulci |
| 2 | Mild atrophy—subtle prominence of the cisternae and sulci for age |
| 3 | Normal brain volume |
| 4 | Mild increased volume but with maintenance of cisternae and sulci |
| 5 | Mild swelling—some loss of cisternae and sulci but not diffuse |
| 6 | Moderate swelling—diffuse involvement with some loss of cisternae and sulci |
| 7 | Severe swelling—loss of all sulci, presence of cisternal effacement, decreased gray/white matter delineation |
| 8 | Severe swelling—loss of all sulci, presence of cisternal effacement, presence of uncal herniation, loss of gray/white matter delineation |

## 3 Methods

Our goal was to develop an automated approach using statistical modeling to predict the BV score given the acquired images. In the following sections, we first describe a preprocessing procedure for the T1 and T2 scans to reduce the effect of image artifacts and to identify brain tissue. We then motivate and develop biomarkers of the three primary assessments used to determine severity of brain swelling.

### 3.1 Pre-processing

#### 3.1.1 Notation

For participant *i* and modality *τ* ∈ {1, 2} corresponding to T1 and T2, a brain image consists of the voxel vector x*_i_* = {1,…, *V_i_*} where *V_i_* is the total number of voxels in that image. At any voxel *x* ∈ x*_i_*, the intensity *v_iτ_* (*x*) defines a function from the integers to the real numbers. By evaluating *v_iτ_* (*x*) at all voxels, we obtain the vector **v***_iτ_*(*x*), which is collectively referred to as the *image*.

#### 3.1.2 N4 Bias Correction

A common artifact of the MRI acquisition process is intensity inhomogeneity or bias, wherein the intensities vary in a gradient over the entire image. Since this can affect the validity of subsequent analyses that assume that a given tissue has the same intensity regardless of its position in the image, bias correction is a common pre-processing step in neuroimaging studies (Sled et al., 1998; Vovk et al., 2007). We corrected all images in our sample using N4 bias correction (Tustison et al., 2010), which assumes a multiplicative bias model for the observed image:

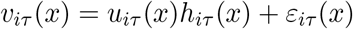

for participant *i* and modality *τ*, such that *u_iτ_* is the true image, *h_iτ_* is a smooth bias field, and *ε_iτ_* is Gaussian noise that is independent of *u_iτ_* (Web Figure 1). To render the model identifiable, it is common to assume a log-additive model where the noise is removable,

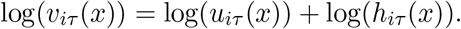

**Figure 1:**
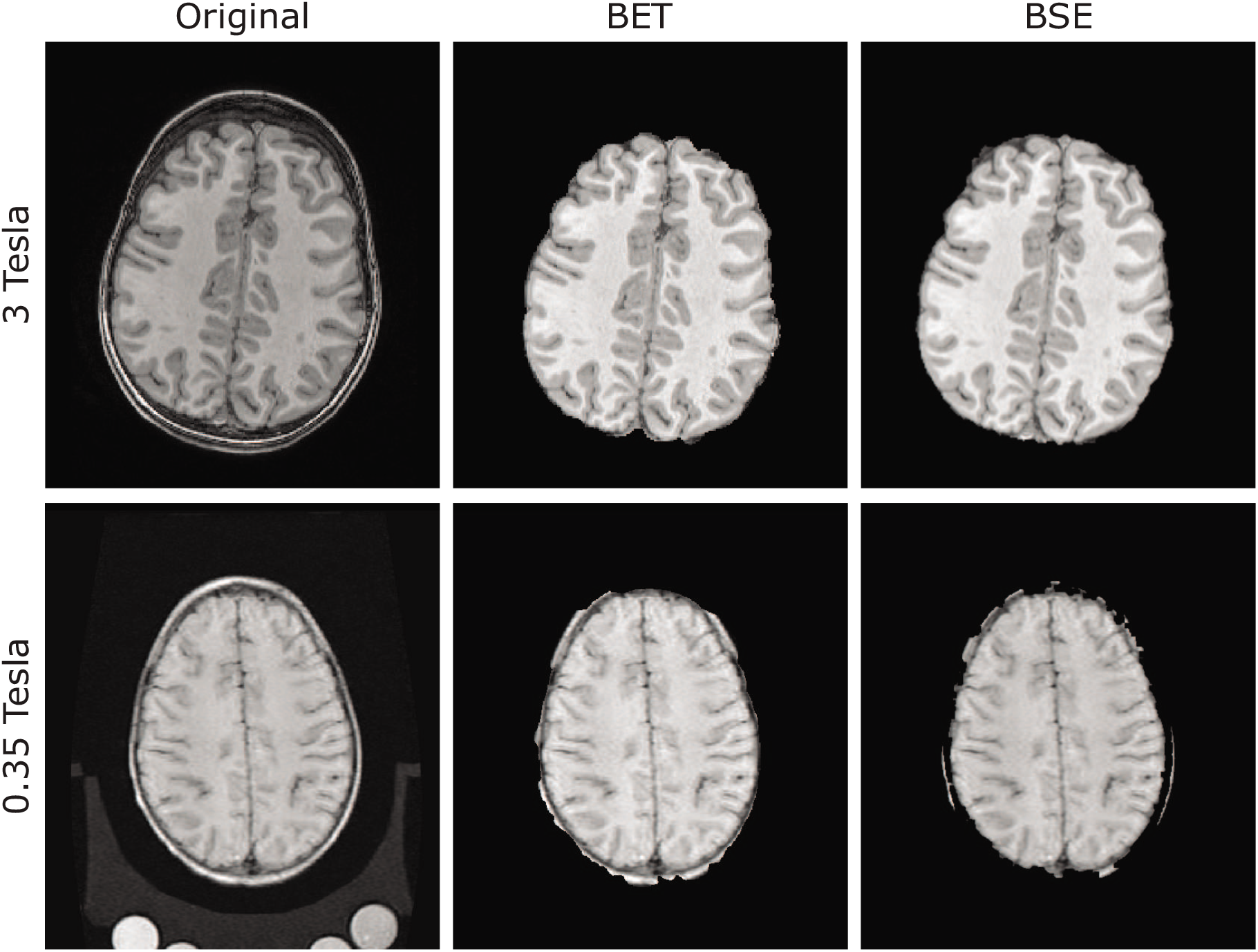
Surface-based brain segmentation methods such as Brain Extraction Toolbox (BET) and Brain Surface Extractor (BSE) perform well on higher resolution compared to lower resolution images. (Top row) BET and BSE segmentation on a 3T image from the Philadelphia Neurodevelopmental Cohort (Satterthwaite et al., 2016). (Bottom row) BET and BSE segmentation on a 0.35T image from our sample. Notably, the 0.35T segmented image still contains much of the skull, which is completely removed in the 3T segmented image.

To estimate the corrected image, we assume that the true image log(*u_iτ_*(*x*)) and log(*h_iτ_* (*x*)) are uncorrelated or independent random variables, that the distribution of log(*h_iτ_*(*x*)) is Gaussian, and that the distribution of log(*u_iτ_*(*x*)) can be reached by deconvolving the distribution of the observed image log(*v_iτ_* (*x*)) from the distribution of log(*h_iτ_*(x)). The bias field *h_iτ_*(*x*) is approximated by the following iterative process: at step 0,

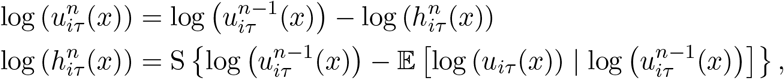

with iteration until the estimated bias field converges. The operator S{·} is a B-spline operator ensuring that the resulting bias field is smooth. The expectation is based on the estimated distribution of log(*u_iτ_*(*x*)) mentioned above; the full derivation can be found in (Tustison et al., 2010).

#### 3.1.3 Brain Segmentation

Because the skull and other non-brain tissues contain noisy and extraneous information, we performed tissue segmentation, where voxels corresponding to a tissue of interest are identified. In the case of brain segmentation, we define the class assignment for voxel *x* and participant *i*:

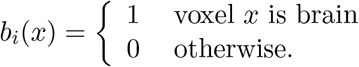

The voxelwise evaluation of *b_i_*(*x*) at *x* ∈ {1,…, *V_i_*} yields the person-specific brain mask *b_i_*, a binary vector of length *V_i_* where the *i*th entry corresponds to the classification of the ith voxel as either brain or non-brain.

We first assessed the performance of two popular surface-based brain segmentation tools, the Brain Extraction Tool (BET) from the FMRIB Software Library (FSL) (Smith, 2002), and the Brain Surface Extractor (BSE) from BrainSuite (Sandor and Leahy, 1997). Both BET and BSE are methods designed for high-resolution images (Boesen et al., 2004; Shi et al., 2012).

While both programs performed well on the higher-resolution 3T images, their performance deteriorated on the low-resolution images in our sample, as the larger voxel size precluded a clear separation between brain and skull (Figure 1).

In response, we appealed to a class of methods that borrow strength from existing “gold standard” segmentations on atlases, which consist of a high-resolution brain image together with its expert-validated brain mask. Our atlas set comprises a sample of 12 participants who were imaged in a high-field 3T scanner as part of the study-specific atlas in the Philadelphia Neurodevelopmental Cohort (Satterthwaite et al., 2016). For these participants, the whole brain was automatically segmented and masks were manually corrected slice by slice, a time-intensive process. The 12 youngest participants in this study cohort (mean age ± sd = 10.06 ± 0.91 years) were chosen to minimize age-related biases that may arise from atlases created from images of patients who are older than the participants in our CM sample.

Atlas-based brain segmentation is typically a two-stage process. First the atlas is continuously deformed to match the target image (that is, the image to be segmented); this deformation is referred to as the registration function. We used symmetric image normalization (SyN) to estimate non-linear registration functions (Avants et al., 2008). Second, the registration function is applied to the atlas’s brain mask, yielding a mask warped into the target image coordinate space indicating where various tissues are located in the target image. To address heterogeneity across participants under study, it is common practice to repeat this process using multiple atlases and brain masks; such methods are referred to as multi-atlas methods (Rohlfing et al., 2004).

The next step is to produce a consensus segmentation of the target in a process called label fusion. We employed a majority voting consensus algorithm (Artaechevarria et al., 2009). At each voxel, the final designation of brain versus non-brain was decided by the majority of warped atlas brain masks at that voxel. Although majority voting has been criticized as overly simplistic, it performed well in our data compared to more advanced label fusion methods (Wang et al., 2013), likely due to lower image quality. Compared to BET and BSE, the multi-atlas method resulted in the most accurate brain segmentation (Web Figure 2), particularly at the inferior and superior extremes. These errors are likely due to the large voxels (i.e., lower resolution) of our data and lack of clear separation between brain and other tissues, which are important to surface-based methods such as BSE and BST. In addition, the limited field of view in most of our images resulted in significant errors from BSE and BST near the vertex (top of the head).

**Figure 2:**
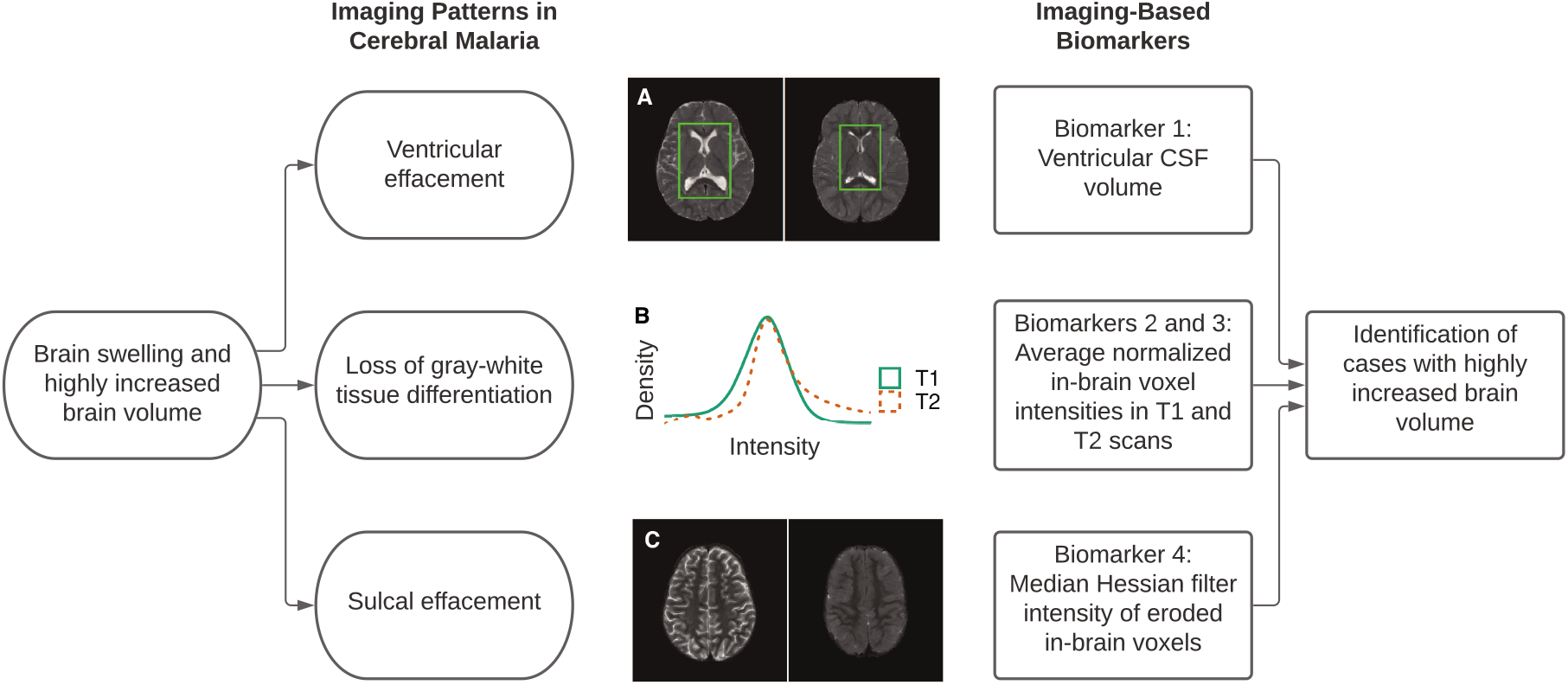
Schematic of the imaging patterns and radiological criteria used to identify highly increased brain swelling on MRI in patients with cerebral malaria. Each characteristic is associated with a corresponding image-based biomarker calculated from the observed T1 and T2 scans from each individual. Panel A: Ventricular CSF volume and BV score. In these T2-weighted images, CSF is shown as bright regions and the ventricles are highlighted by green boxes. (Left) A participant with BV score = 2.5 indicating near-normal brain volume. (Right) A participant with BV score = 7 indicating highly increased brain volume. Panel B: intensity histogram of in-brain voxels in T1 and T2 images. Panel C: Gyral features in the T2-weighted MRI scan and BV score. (Left) A scan showing the sulci as bright for a participant with BV score = 2, indicating near-normal brain volume. (Right) A scan from a participant with BV score = 8 indicating severely increased brain volume. The sulci appear noticeably darker as they no longer contain CSF.

### 3.2 Biomarkers

Based on the radiological criteria for BV scores of 7 and 8 (Table 1), we developed three types of image-based multi-modal biomarkers to quantify a) ventricular volume, b) gray and white matter delineation, and c) sulcal effacement.

#### 3.2.1 Ventricular CSF Volume

We hypothesized that brain swelling would affect the appearance of the CSF-containing cerebral ventricles (Figure 2). Ventricular effacement (which results when the ventricular volume of CSF is diminished), along with sulcal and cisternal effacement, is frequently noted in neuroimaging from patients with non-malarial brain swelling (Ho et al., 2012) and cerebral malaria (Mishra and Newton, 2009). The high signal of CSF on T2-weighted images, and the availability of open-access ventricle atlases, makes the identification of ventricular CSF an appealing candidate for automated tools.

A measure of ventricular CSF requires the identification of both ventricular and CSF voxels in the image. To identify CSF regions, we leveraged a model of the observed intensities to partition voxels into three tissue classes: white matter, gray matter, or CSF. We used FSL FAST (Zhang et al., 2001), a popular approach that assumes that intensities and tissue classes can be modeled by a Gaussian Hidden Markov Random Field (GHMRF). For voxel *x* and person *i*, this assignment can be summarized by a tissue class segmentation function *w_i_*(*x*):

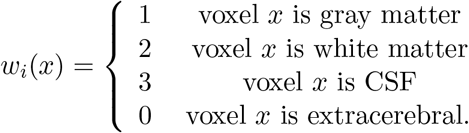

The collective tissue class assignments obtained by evaluating *w_i_*(*x*) at all voxels are denoted **w***_i_*. To estimate **w***_i_* in the GHMRF, both the observed intensities **v***_iτ_* and the true tissue classes **w***_i_* are considered to be random, and the goal is to find the class assignment 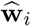 maximizing their joint likelihood

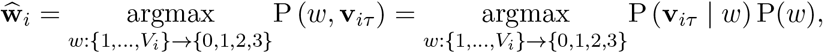

where the conditional distribution of *v_iτ_*|*w* is assumed to be Gaussian, and the marginal distribution of tissue classes w follows a Gibbs distribution.

The conditional Gaussian assumption is standard in brain tissue segmentation methods (Avants et al., 2011; de Boer et al., 2010), though it is often not met in practice, as was the case in our data (Web Figure 4). In particular, Shapiro-Wilks tests showed that the conditional distributions of intensities within each tissue class were not Gaussian; indeed, intensities were often bimodally distributed within the class identified as CSF. Mapping these voxels back to the brain image revealed that one of the peaks corresponded to incorrectly segmented gray and white matter. By limiting our analyses to CSF-labeled voxels within the ventricles (as opposed to whole-brain CSF), the majority of these incorrectly labeled voxels were removed (Web Figure 4). Although the Gaussian distributional assumptions were not met, the resulting segmentation performed well.

Another motivation for choosing ventricular CSF volume over whole-brain CSF volume is that the latter was highly variable among participants, especially at the brain boundary. This is likely due to the brain segmentation performed in the previous step. The ventricular CSF volume yielded a more stable estimate and demonstrated better identification of participants with highly increased BV than either whole-brain CSF volume or ventricular volume alone (Web Figure 3).

**Figure 3:**
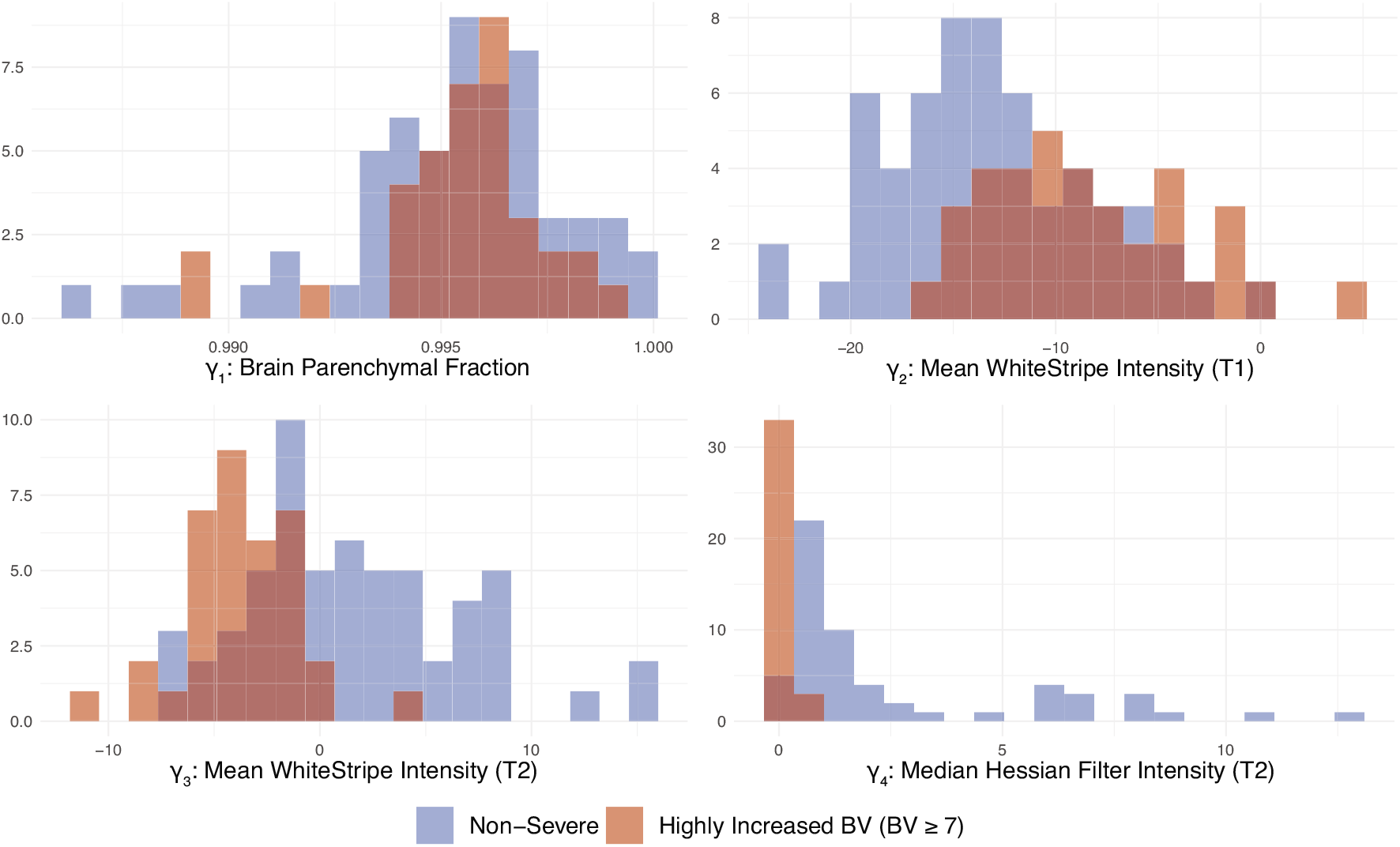
Distribution of derived biomarkers over all participants in the sample and categorized by severe edema status.

To identify voxels corresponding to the ventricles, we obtained adult ventricle atlases from the publicly available OASIS cross-sectional data set (Marcus et al., 2007). Using the same procedure of SyN registration and majority-voting label fusion as in the brain segmentation step, we obtained a ventricle mask for each participant, and calculated the ventricular CSF mask as the intersection of the CSF mask from FSL FAST and the OASIS ventricle mask.

For voxel *x* and participant *i*, we define the ventricular CSF mask function

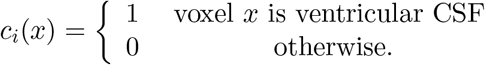

The first image-based biomarker is the brain parenchymal fraction of the T2 scan, or the proportion of non-ventricular CSF voxels to total brain voxels:

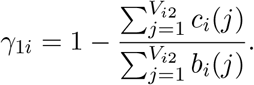

Higher values of BPF represent a lower proportion of ventricular CSF volume and thus higher levels of brain swelling (Figure 3).

#### 3.2.2 Gray and White Matter Intensity

To translate the loss of gray and white matter delineation into a function of the observed image, it is necessary to normalize voxel intensities within each modality so that participants can be compared. In MRI scans, voxel intensities are measured in non-standardized units with no biological meaning; there can be significant variation in the appearance of these images even within the same participant (Nyul et al., 2000). To normalize the intensities, we applied a linear scaling of the image intensities based on normal-appearing white matter (NAWM) using the WhiteStripe technique (Shinohara et al., 2014).

WhiteStripe intensity normalization depends on the identification of white matter voxels in the brain image. In theory, each brain image contains the same structures (i.e., white matter, gray matter, CSF) and we assume that magnetic resonance imaging produces different intensity distributions for these tissues (Tohka et al., 2007). The overall distribution of intensities can be modeled as a mixture distribution, and the number of components is typically determined *a priori*s based on our knowledge of the image contents (Web Figure 5). In practice, Gaussian mixture models are popular in medical imaging due to their simplicity and speed (Balafar, 2012; Shattuck et al., 2001).

We rely on this working assumption in order to easily identify the white matter voxels for WhiteStripe. For participant *i* and modality *τ*, the observed intensities **v***_iτ_* are assumed to follow a mixture distribution with *K* components. That is, the probability density 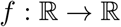 is a function of intensity value v that decomposes as

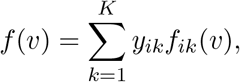

where the 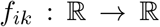 are person-specific probability density functions and the weights *y_ik_* sum to 1. We expect that each individual’s brain image has common structures whose distributions can be mapped across participants, so we make the working assumption that there exists a transformation *f_ik_*(*v*) → *g_k_*(*v*) so that the resulting intensity distribution is not person-specific:

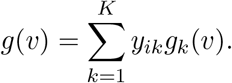

This working assumption allows us to compare intensity distributions from any pair of individuals (Web Figure 6). The white stripe of NAWM is found by smoothing the empirical intensity histogram with a penalized spline (Ruppert et al., 2003) and identifying the peak corresponding to white matter (in T1 sequences this is the rightmost or highest-intensity peak; in T2 sequences this is the overall mode). The interval around this peak, whose width may be adjusted by tuning parameters, is the white stripe. Every voxel intensity within the brain is then linearly scaled by the mode 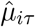 and trimmed standard deviation 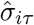 of intensities within the white stripe:

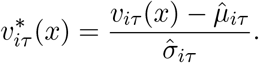

Letting *B_i_* equal the total number of brain voxels for participant *i*, we calculated the second and third biomarkers (Figure 3) using the T1 and T2 images by taking the average normalized intensity after WhiteStripe within the brain voxels only (as determined by the brain segmentation mask in the previous section):

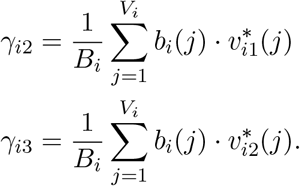

#### 3.2.3 Sulcal Effacement

Finally, we considered that sulcal effacement, which is associated with BV scores of 6 to 8, might be extracted from the MRI by detecting gyri, or ridges, of the cerebral cortex (Figure 2). To do so, we used a filter on the Hessian matrix of each MRI image (Frangi et al., 1998).

In a 3-dimensional image, the Hessian matrix 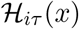 at voxel *x* contains information about the local curvature around *x* for participant *i* and modality *τ*. Typically, 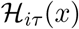 is calculated by convolving a neighborhood of *x* with derivatives of a Gaussian kernel. The three eigenvalues of 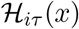 with smallest magnitudes 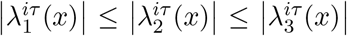 have a geometric interpretation: gyral or planar structures correspond to small values of 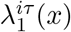 and 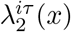, and a high value of 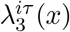 (Frangi et al., 1998). Tubular and spherical structures, on the other hand, are associated with different patterns in these eigenvalues, so the following dissimilarity measures are required to identify gyral features:

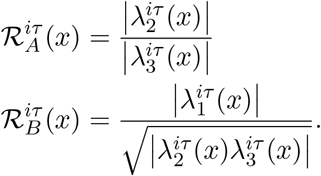

The resulting Hessian filter is defined as

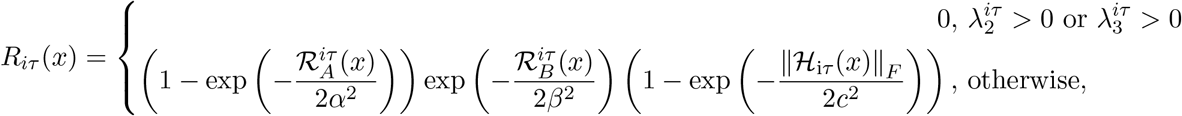

where ║·║*_F_* is the Frobenius norm and *α, β*, and *c* are parameters tuning the importance of 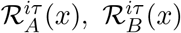, and 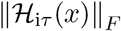 respectively. By calculating *R_iτ_*(*x*) at each voxel, we produced a probability-like map highlighting gyral features.

We applied the Hessian filter implemented in ITK-SNAP’s Convert3D tool (Yushkevich et al., 2006) to the T2 images, as they showed the best contrast between CSF and brain tissue. Due to limited field of view and the quality of brain segmentation around the top and bottom of the brain, the Hessian filter was only calculated in MRI slices taken from central portions of the cerebral hemispheres. The middle portion was defined by removing the inferior and superior 3 slices (i.e., the top and bottom) of the axial sequence of each T2 image, as well as any voxels neighboring extracranial tissue (Web Figure 7).

Similar to the brain mask *b_i_*(*x*) defined in the previous section, the eroded brain mask is defined as

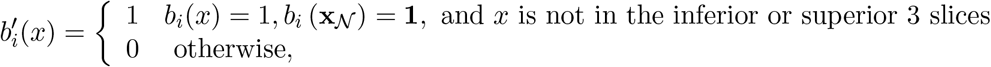

where 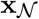 is the set of voxels adjacent to *x*.

We defined the final biomarker (Figure 3) as the median of the Hessian filter intensities in the T2 image limited to eroded brain voxels 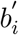:

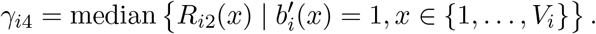

The median was chosen as the distributions of the Hessian filter intensities within each image were highly right skewed; this resulted in a more conservative and robust characterization of the difference between severe and non-severe cases.

## 4 Prediction of Highly Increased Brain Volume in Cerebral Malaria

### 4.1 Model

To predict highly increased brain volume (a median BV score of 7 or higher), we defined the binary outcome variable for participant *i* and median BV score threshold *k* as the indicator variable 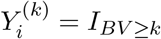. The main outcome of interest in our study was 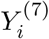. Together with the biomarkers introduced in the previous section, we formed the multivariate logistic regression model

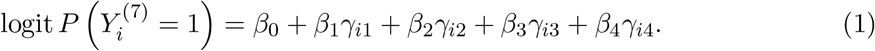

All statistical analyses were performed in R version 3.6.1 (R Core Team, 2019). The estimated coefficients 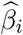 fit on the training set are shown in Table 2. Participants with a BV score of 7-8 were associated with a lower median Hessian filter (*p* < 0.01), signifying fewer prominent sulci in the brain. This was the only coefficient found to be significantly different from zero, consistent with radiologists’ reports that sulcal effacement was the most important factor in determining a higher BV score. Both the intercept term and the coefficient of BPF had high standard errors, suggesting near-complete data separation. Consequently, we did not interpret those coefficients.

**Table 2:** Estimated coefficients in the prediction model of highly increased BV using the training data (*n* = 48). The Hessian intensity biomarker (*γ*_.4_) was the sole significant covariate, supporting clinical observations that sulcal effacement was the most important factor in determining cases scoring 7 or higher.

|  | *Dependent variable:* |
| --- | --- |
| | BV Score $\geq 7$ ($Y^{(7)}$) |
| $\gamma_1$ (Brain Parenchymal Fraction) | -171.58<br>(-702.45, 359.29) |
| $\gamma_2$ (Mean WhiteStripe Intensity - T1) | 0.08<br>(-0.17, 0.33) |
| $\gamma_3$ (Mean WhiteStripe Intensity - T2) | -0.11<br>(-0.66, 0.43) |
| $\gamma_4$ (Median Hessian Intensity - T2) | -8.12***<br>(-14.17, -2.06) |
| Constant | 174.19<br>(-354.58, 702.95) |
| Observations | 48 |
| Log Likelihood | -10.68 |
| Akaike Inf. Crit. | 31.35 |
| *Note:* | *p<0.1; **p<0.05; ***p<0.01 |

### 4.2 Prediction Accuracy

In addition to model 1, we performed several sensitivity analyses. The biomarkers *γ_.j_* for *j* ∈ {1,…, 4} were developed to identify images with BV ≥ 7; we considered an additional set of models with outcomes 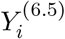 and 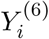, where the threshold for determining severity was relaxed. Because *γ*_*i*4_, the measure of sulcal effacement, was both the most statistically and clinically important predictor of highly increased brain volume cases, we also considered models with *γ*_*i*4_ alone (“Hessian Only”) as opposed to the full set of covariates (“All Biomarkers”). In total, we examined 5 alternate models in addition to the main model, which are summarized in Table 3.

**Table 3:** Model prediction is highly accurate regardless of BV threshold. All logistic models were trained on the training set, and accuracy was evaluated on both the training set and the withheld test set. Area under the curve (AUC) and 95% confidence interval for logistic regression models with binary outcomes 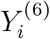 (BV ≥ 6), 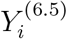 (BV ≥ 6.5), and 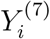 (BV ≥ 7); and either the full set of covariates in Model 1 (“All Biomarkers”) or with the sulcal effacement biomarker (“Hessian Only”). The primary model in 1 is shown in bold.

| Outcome | Covariates | Train AUC | Train AUC<br>(95% CI) | Test AUC | Test AUC<br>(95%CI) |
| --- | --- | --- | --- | --- | --- |
| **$BV \geq 7$** | **All Biomarkers** | **0.97** | **[0.93,1.00]** | **0.95** | **[0.89,1.00]** |
| $BV \geq 7$ | Hessian Only | 0.97 | [0.92,1.00] | 0.96 | [0.90,1.00] |
| $BV \geq 6.5$ | All Biomarkers | 0.97 | [0.93,1.00] | 0.92 | [0.83,1.00] |
| $BV \geq 6.5$ | Hessian Only | 0.97 | [0.92,1.00] | 0.93 | [0.84,1.00] |
| $BV \geq 6$ | All Biomarkers | 0.96 | [0.91,1.00] | 0.97 | [0.93,1.00] |
| $BV \geq 6$ | Hessian Only | 0.81 | [0.67,0.95] | 0.90 | [0.78,1.00] |

We assessed prediction accuracy using area under the Receiver Operating Characteristic curve (AUC), which considers the sensitivity and specificity across different thresholds of the predicted outcome. For all models, the AUC was high, reaching 0.81 to 0.97 in the training set, and 0.90 to 0.97 in the testing set. Models with outcome 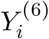 received a lower AUC than those with outcome 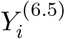 and 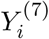, although the 95% confidence intervals for all models intersect. This finding—in conjunction with clinical observations—suggests that patients with a BV score of 6 represent borderline cases and are therefore graded with the most uncertainty.

We compared models with *γ*_*i*4_ as the sole covariate to the full model using a likelihood ratio test, and found that the reduced model performed similarly when the outcome was 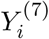 (*p* > 0.3 in both training and test sets), moderately when the outcome was 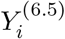 (*p* < 0.01 in the testing set only), and poorly when the outcome was 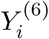 (*p* < 0.01 in both training and test sets). In other words, the sulcal effacement biomarker was sufficient to classify cases scoring a 7-8, but all 3 biomarkers were needed to accurately classify cases with BV scores of 6-8. This suggests that the measures of ventricular CSF and gray-to-white matter differentiation, while less relevant for predicting cases scoring 7 or higher, may be useful in differentiating cases that assigned a score of 5 or 6.

In practical settings, a threshold on the predicted probability in model (1) is required to determine which individuals are predicted as severe or non-severe. While this can be done using cross-validation, we also propose the following method to identify an “ideal” threshold which maximizes sensitivity and specificity in that order, with the restriction that specificity be greater than 0.9 (Web Table 3). Due to the discrete nature of the observations, the ideal threshold is not necessarily unique. For outcome 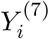, the sensitivity was 0.78 and specificity was 0.96 using the full set of covariates; the sensitivity was 0.94 and specificity was 0.93 using just the sulcal effacement covariate. Together, these support the previous conclusion that sulcal effacement is the most important predictor for BV scores of 7 or higher. We have also identified a decision rule to distinguish severe and non-severe BV with low rates of false negatives and false positives. Indeed, a wide interval of acceptable thresholds, ranging from around 0.3 to 0.8, will yield acceptable values (>75%) of sensitivity, specificity, positive predictive value (PPV), and negative predictive value (NPV) (Web Figure 10). This suggests that, in clinical settings, the ultimate determination of severe or non-severe cases will have good accuracy that is robust to the choice of threshold (Web Table 4).

**Table 4:** Nested cross-validation reveals that our findings are not dependent on the initial split of training and test data. Averaged over 5 splits, the mean and 95% confidence interval of sensitivity, specificity, Youden’s *J* (informedness), negative predictive value (NPV), and positive predictive value (PPV) are shown for logistic regression models with binary outcomes 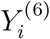 (BV ≥ 6), 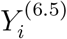 (BV ≥ 6.5), and 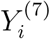 (BV ≥ 7); and either the full set of covariates in Model (1) (“Full Set”) or with the sulcal effacement biomarker (“Hessian Only”). The primary model in (1) is shown in bold.

| Outcome | Covariates | Sensitivity | Specificity | Youden’s<br>$J$ | NPV | PPV |
| --- | --- | --- | --- | --- | --- | --- |
| **$Y_i^{(7)}$** | **Full Set** | **0.89**<br>**(0.75,1)** | **0.92**<br>**(0.68,1)** | **0.82**<br>**(0.66,0.97)** | **0.85**<br>**(0.65,1)** | **0.95**<br>**(0.81,1)** |
| $Y_i^{(7)}$ | Hessian<br>Only | 0.91<br>(0.79,1) | 0.92<br>(0.76,1) | 0.83<br>(0.57,1) | 0.86<br>(0.68,1) | 0.94<br>(0.84,1) |
| $Y_i^{(6.5)}$ | Full Set | 0.9<br>(0.75,1) | 0.83<br>(0.48,1) | 0.73<br>(0.24,1) | 0.82<br>(0.56,1) | 0.9<br>(0.68,1) |
| $Y_i^{(6.5)}$ | Hessian<br>Only | 0.88<br>(0.73,1) | 0.91<br>(0.67,1) | 0.79<br>(0.7,0.89) | 0.82<br>(0.62,1) | 0.93<br>(0.67,1) |
| $Y_i^{(6)}$ | Full Set | 0.81<br>(0.69,0.93) | 0.9<br>(0.61,1) | 0.71<br>(0.4,1) | 0.84<br>(0.64,1) | 0.83<br>(0.36,1) |
| $Y_i^{(6)}$ | Hessian<br>Only | 0.23<br>(0,0.54) | 0.87<br>(0.67,1) | 0.1<br>(0,0.29) | 0.58<br>(0.41,0.74) | 0.65<br>(0.18,1) |

### 4.3 Resampling Validation

Although there were no known systematic clinical differences in the training and test set collection, there was a difference in the number of raters. To confirm that our results were not dependent on the initial split, we combined the training and test datasets and assessed prediction accuracy using nested cross-validation. In the inner loop, 4-fold cross-validation was used to fit the model parameters and determine the optimal cutoff; in the outer loop, 5-fold cross-validation was used to assess the trained model on a validation set.

In the resampling scheme, we observed the same pattern in prediction performance: classification accuracy remained high in nearly all models (Table 4). We again found that the Hessian Only model performed as well as the full model for 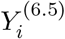 and 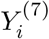, but notably worse than the full model for 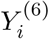. Together, these results show that our derived biomarkers and model are able to accurately and robustly differentiate between participants with highly increased brain volume (BV scores of 7 or 8) from those who do not. The measurement of sulcal effacement *γ*_*i*4_ was found to be the most important factor to identify cases with BV ≥ 7 with high sensitivity, specificity, PPV, and NPV at selected thresholds of the predicted outcome (Web Table 3). Meanwhile, measures of ventricular CSF volume and gray and white matter differentiation (*γ*_*i*1_, *γ*_*i*2_, and *γ*_*i*3_) provided more value in identifying “borderline” cases. This finding aligns with radiologists’ reports that images with scores ranging from 5 to 6.5 relied on heavily subjective calls. To score such cases in a clinical setting would usually require a consensus among several radiologists; we conjecture that this uncertainty is reflected in the poorer performance of the model in classifying these cases.

### 4.4 External Validation

Next, we tested the performance of our model on external data from an ongoing study of brain swelling in CM. The external data included 90 new participants aged 6 months to 11 years old (mean age ± sd = 4.9 ± 2.4 years). All participants had both axial T1 and T2 scans, were imaged on the same scanner as the participants in our original data, and were scored by two of the same radiologists as the original data. (Imaging protocols were similar between the new data and original data, with details available upon request.)

After fitting model (1) on the complete set of original data (*n* = 94), we assessed performance on this new external data. The resulting models achieved good prediction accuracy (Web Table 5), albeit slightly reduced compared to the original data. For models predicting BV scores of 7-8 using either the full set of covariates and the sulcal effacement biomarker, the AUC in the external dataset was 0.88 (95% CI: 0.80 to 0.96). Overall, these findings demonstrate that our method is applicable to other data sets of children with CM.

### 4.5 Linear Modelling

We investigated if it was possible to predict the BV score directly, rather than as a binary outcome. Using multivariate linear regression combined with nested cross-validation, we found that a higher BV score (worse swelling) was significantly associated with a higher *γ*_*i*2_, lower *γ*_*i*3_, and lower *γ*_*i*4_ (Web Table 1), aligning with our expectations and the observed distribution of these biomarkers in highly increased and non-highly increased cases (Figure 3). The prediction error was adequate (RMSE = 1.13), though the linear model performed less well in predicting highly increased cases: after thresholding the predicted score at 7, the predicted binary outcome had an average sensitivity of 0.95, a specificity of 0.35, and a Youden’s *J* of 0.3 (Web Table 2). Compared to the logistic model, the lower specificity suggests that the linear model is more likely to produce false positives of highly increased brain volume cases.

### 4.6 Alternate Biomarker Specifications

We also considered using different summary statistics in formulating biomarkers *γ*_*i*1_, *γ*_*i*2_, and *γ*_*i*3_. We originally used the average WhiteStripe intensities and median Hessian intensities, but also assessed the performance of models using the median WhiteStripe intensities and average Hessian intensities, and all varying combinations thereof. Overall, all models performed comparably (Web Figure 8), suggesting that our method is relatively robust to the choice of summary statistic. This is likely due to the high correlations between the mean and median for each variable (Web Figure 9).

### 4.7 Comparison to Human Raters

Finally, we compared the performance of the model (1) prediction to that of the human raters in predicting cases with BV ≥ 7. The performance of each human rater was measured against the “ground truth” defined by the consensus of the remaining raters. Because the test set and training set had different sets of raters, we repeated this analysis within each set. We also considered the model with only the Hessian predictor as well as the full set of covariates. In all cases, we found that the sensitivity and specificity of the model was comparable to that of the human raters (Figure 4); in particular, our proposed method out-performed around half of the human raters in detecting true negative cases.

**Figure 4:**
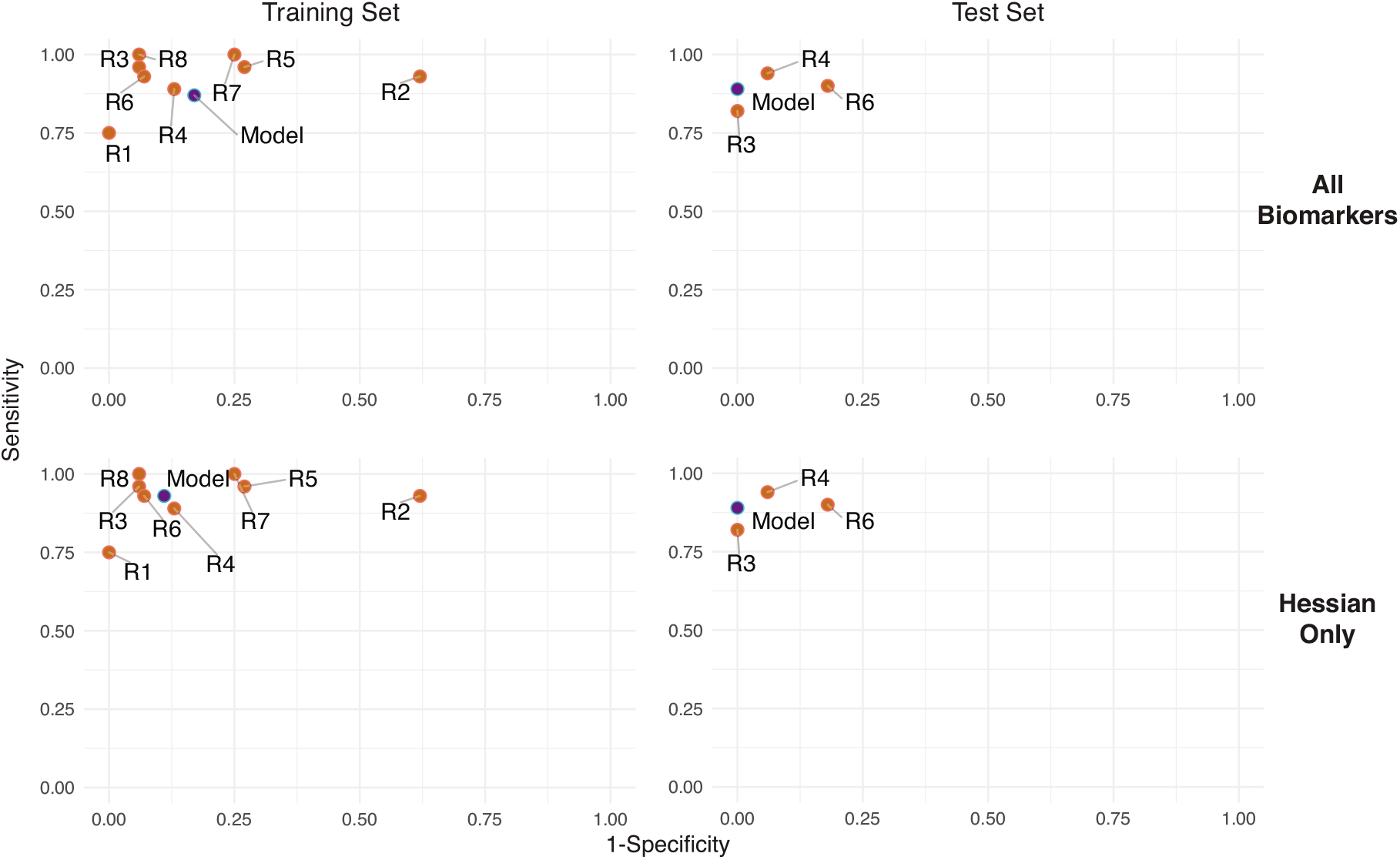
Compared to the human raters (R1 through R8; orange dots), our automated method (blue dot) performs comparably in identifying patients with BV ≥ 7. Here, we assess prediction performance using sensitivity and 1-specificity (false positive rate), where points in the upper left quadrant correspond to better performance. The automated prediction was fit on either the training or test set, and evaluated for sensitivity and specificity on the other set. The accuracy of human raters was determined against the median of the other raters. Whether the model was fit on all four imaging biomarkers (top row) or just the Hessian intensity biomarker (bottom row), the automated prediction performed better than around half of the human raters in terms of specificity, and performed worse in terms of sensitivity.

## 5 Discussion

We developed a method for the statistical image analysis of low-resolution, noisy brain MRI, after determining that a popular method developed for images obtained on high-field MRI scans did not perform well on the images in our sample. Our method involved creating and validating a multi-atlas, integrative framework to automate the radiological assessment of brain volume, a biomarker strongly associated with death in children with CM. Our logistic classification model is parsimonious, aligns with clinical observations, and has high predictive accuracy based on multiple performance metrics (Adhikari et al., 2021). This excellent performance extends to previously unseen data from new study participants.

The automated pipeline relies only on R and other open-source programs; all code and prediction model coefficients will be made available on GitHub. High-resolution atlases for pediatric brains (Fonov et al., 2011) and adult ventricles (Marcus et al., 2007) are also publicly available, so that other researchers can easily implement the pipeline. An attractive feature of this program is that it requires only a participant’s T1- and T2-weighted MRI sequences as input in order to predict highly increased BV. We hope that the implementation of these findings in low-resource environments might help to address both the shortage of radiologists for manual MRI interpretation, as well as the challenge of interpreting images from low field MRI.

Our results provide insight into how radiologists score brain edema on MRI, generally supporting the stated importance of sulcal effacement over ventricular size and gray-white matter delineation. The superior performance of the sulcal effacement biomarker suggests that higher scores of brain edema were predominantly derived from this MRI finding, even though other features such as loss of gray-to-white matter delineation are also prescribed features for images assigned BV scores of 7 to 8. Future efforts to provide Hessian filter images or sulcal effacement scores might assist individual radiologists in producing more consistent gradings of brain volume. We intend to apply these methods to follow-up analyses of brain MRI scans obtained with even lower field (< 0.1 T) MRI scanners that are now being introduced into clinical use in both high and low-resource settings (Sheth et al., 2020), including to evaluate CM.

Another potential application of our pipeline is to provide image-based decision support. For instance, we demonstrated that the automated prediction performs as well as the average human rater in our sample. This could be useful in situations where a consensus interpretation is required, such as borderline cases which are suspected to score between 5 to 6.5. Replacing one of the human raters with the model rating (Botwe et al., 2021) could also curb the logistical complication of coordinating multiple radiologists, since only a few are on call at any one time.

A limitation of our approach is that logistic regression only accommodates binary outcomes (highly increased and not highly increased BV scores). Linear models which predicted BV score directly have higher false positive rates (Web Table 2), suggesting that more information or more sophisticated models may be needed to predict the ordinal score. Future analyses could apply our pipeline to predict disease outcome: Kampondeni et al. (2018) showed that global CSF volume was the best predictor of prognosis in patients with CM. However, the accuracy of global CSF measurements in our current pipeline is limited by the quality of segmentation at the edge of the brain. Recent developments in deep learning methods for brain segmentation (Ronneberger et al., 2015) could address this issue, although such procedures may require larger sample sizes, with manual delineations of brain tissue, than are currently available.

In summary, we introduced and validated a biologically and statistically principled method of biomarker development using images from low field strength MRIs, including images with artifacts. These strategies, which involve borrowing strength from publicly available high-resolution data whenever possible, and considering aggregate statistics that are more robust to extreme values, can be applied to any study of low-resolution brain images. The principles behind the tools introduced in this study are also broadly applicable to the design of new techniques that automate existing, clinically validated tasks.

## References

Adhikari, S., Normand, S.-L., Bloom, J., Shahian, D., and Rose, S. (2021). Revisiting performance metrics for prediction with rare outcomes. Statistical Methods in Medical Research 30, 2352–2366.

Artaechevarria, X., Munoz-Barrutia, A., and de Solorzano, C. O. (2009). Combination strategies in multi-atlas image segmentation: Application to brain MR data. IEEE Transactions on Medical Imaging 28, 1266–1277.

Avants, B., Epstein, C., Grossman, M., and Gee, J. (2008). Symmetric diffeomorphic image registration with cross-correlation: Evaluating automated labeling of elderly and neurodegenerative brain. Medical Image Analysis 12, 26–41.

Avants, B. B., Tustison, N. J., Wu, J., Cook, P. A., and Gee, J. C. (2011). An open source multivariate framework for n-tissue segmentation with evaluation on public data. Neuroinformatics 9, 381–400.

Balafar, M. A. (2012). Gaussian mixture model based segmentation methods for brain MRI images. Artificial Intelligence Review 41, 429–439.

Bhat, S. S., Fernandes, T. T., Poojar, P., Ferreira, M. S., Rao, P. C., Hanumantharaju, M. C., Ogbole, G., Nunes, R. G., and Geethanath, S. (2020). Low-field MRI of stroke: Challenges and opportunities. Journal of Magnetic Resonance Imaging.

Boesen, K., Rehm, K., Schaper, K., Stoltzner, S., Woods, R., Lüders, E., and Rottenberg, D. (2004). Quantitative comparison of four brain extraction algorithms. NeuroImage 22, 1255–1261.

de Boer, R., Vrooman, H. A., Ikram, M. A., Vernooij, M. W., Breteler, M. M., van der Lugt, A., and Niessen, W. J. (2010). Accuracy and reproducibility study of automatic MRI brain tissue segmentation methods. NeuroImage 51, 1047–1056.

Dondorp, A. M., Fanello, C. I., Hendriksen, I. C., Gomes, E., Seni, A., Chhaganlal, K. D., and Others (2010). Artesunate versus quinine in the treatment of severe falciparum malaria in african children (AQUAMAT): an open-label, randomised trial. The Lancet 376, 1647–1657.

Fonov, V., Evans, A. C., Botteron, K., Almli, C. R., McKinstry, R. C., and Collins, D. L. (2011). Unbiased average age-appropriate atlases for pediatric studies. NeuroImage 54, 313–327.

Frangi, A. F., Niessen, W. J., Vincken, K. L., and Viergever, M. A. (1998). Multiscale vessel enhancement filtering. In International conference on medical image computing and computer-assisted intervention, pages 130–137. Springer.

Gordillo, N., Montseny, E., and Sobrevilla, P. (2013). State of the art survey on MRI brain tumor segmentation. Magnetic Resonance Imaging 31, 1426–1438.

Ho, M.-L., Rojas, R., and Eisenberg, R. L. (2012). Cerebral edema. American Journal of Roentgenology 199, W258–W273.

Idro, R., Marsh, K., John, C. C., and Newton, C. R. J. (2010). Cerebral malaria: Mechanisms of brain injury and strategies for improved neurocognitive outcome. Pediatric Research 68, 267–274.

Kampondeni, S., Seydel, K. B., Zhang, B., Small, D. S., Birbeck, G. L., Hammond, C. A., Chilingulo, C., Taylor, T. E., and Potchen, M. J. (2020). Amount of brain edema correlates with neurologic recovery in pediatric cerebral malaria. The Pediatric Infectious Disease Journal 39, 277–282.

Kampondeni, S. D., Birbeck, G. L., Seydel, K. B., Beare, N. A., Glover, S. J., Hammond, C. A., Chilingulo, C. A., Taylor, T. E., and Potchen, M. J. (2018). Noninvasive measures of brain edema predict outcome in pediatric cerebral malaria. Surgical Neurology International 9, 53.

Kampondeni, S. D., Taylor, T. E., Birbeck, G. L., Potchen, M. J., Seydel, K. B., Beare, N. A. V., and Glover, S. J. (2013). MRI findings in a cohort of brain injured survivors of pediatric cerebral malaria. The American Journal of Tropical Medicine and Hygiene 88, 542–546.

Latourette, M. T., Siebert, J. E., Barto, R. J., Marable, K. L., Muyepa, A., Hammond, C. A., Potchen, M. J., Kampondeni, S. D., and Taylor, T. E. (2010). Magnetic resonance imaging research in sub-saharan africa: Challenges and satellite-based networking implementation. Journal of Digital Imaging 24, 729–738.

Looareesuwan, S., Brittenham, G. M., Laothamatas, J., and Brown, T. R. (2009). Cerebral malaria: A new way forward with magnetic resonance imaging (MRI). The American Journal of Tropical Medicine and Hygiene 81, 545–547.

Luzolo, A. L. and Ngoyi, D. M. (2019). Cerebral malaria. Brain Research Bulletin 145, 53–58.

Marcus, D. S., Wang, T. H., Parker, J., Csernansky, J. G., Morris, J. C., and Buckner, R. L. (2007). Open access series of imaging studies (OASIS): Cross-sectional MRI data in young, middle aged, nondemented, and demented older adults. Journal of Cognitive Neuroscience 19, 1498–1507.

Marques, J. P., Simonis, F. F., and Webb, A. G. (2019). Low-field MRI: An MR physics perspective. Journal of Magnetic Resonance Imaging 49, 1528–1542.

Mishra, S. K. and Newton, C. R. J. C. (2009). Diagnosis and management of the neurological complications of falciparum malaria. Nature Reviews Neurology 5, 189–198.

Mohsen, H., El-Dahshan, E.-S. A., El-Horbaty, E.-S. M., and Salem, A.-B. M. (2018). Classification using deep learning neural networks for brain tumors. Future Computing and Informatics Journal 3, 68–71.

Mollura, D. J., Culp, M. P., Pollack, E., Battino, G., Scheel, J. R., Mango, V. L., Elahi, A., Schweitzer, A., and Dako, F. (2020). Artificial intelligence in low- And middle-income countries: Innovating global health radiology. Radiology 297, 513–520.

Nieuwenhuis, M., Schnack, H. G., van Haren, N. E., Lappin, J., Morgan, C., Reinders, A. A., and Others (2017). Multi-center MRI prediction models: Predicting sex and illness course in first episode psychosis patients. NeuroImage 145, 246–253.

Nyul, L., Udupa, J., and Zhang, X. (2000). New variants of a method of MRI scale standardization. IEEE Transactions on Medical Imaging 19, 143–150.

Obungoloch, J., Harper, J. R., Consevage, S., Savukov, I. M., Neuberger, T., Tadigadapa, S., and Schiff, S. J. (2018). Design of a sustainable prepolarizing magnetic resonance imaging system for infant hydrocephalus. Magnetic Resonance Materials in Physics, Biology and Medicine 31, 665–676.

Ogbole, G. I., Adeyomoye, A. O., Badu-Peprah, A., Mensah, Y., and Nzeh, D. A. (2018). Survey of magnetic resonance imaging availability in west africa. Pan African Medical Journal 30,.

Potchen, M., Kampondeni, S., Seydel, K., Birbeck, G., Hammond, C., Bradley, W., DeMarco, J., Glover, S., Ugorji, J., Latourette, M., Siebert, J., Molyneux, M., and Taylor, T. (2012). Acute brain MRI findings in 120 malawian children with cerebral malaria: New insights into an ancient disease. American Journal of Neuroradiology 33, 1740–1746.

Potchen, M. J., Kampondeni, S. D., Ibrahim, K., Bonner, J., Seydel, K. B., Taylor, T. E., and Birbeck, G. L. (2013). NeuroInterp: A method for facilitating neuroimaging research on cerebral malaria. Neurology 81, 585–588.

Potchen, M. J., Milner, D. A., Glover, S. J., Kampondeni, S. D., Taylor, T. E., Lishimpi, K., Utriainen, D., Mwenechanya, M., Hammond, C. A., Sinyangwe, S. S., Seydel, K. B., Haacke, E. M., Birbeck, G. L., and Zeli, E. (2018). 1.5 tesla magnetic resonance imaging to investigate potential etiologies of brain swelling in pediatric cerebral malaria. The American Journal of Tropical Medicine and Hygiene 98, 497–504.

R Core Team (2019). R: A Language and Environment for Statistical Computing. R Foundation for Statistical Computing, Vienna, Austria.

Rohlfing, T., Brandt, R., Menzel, R., and Maurer, C. R. (2004). Evaluation of atlas selection strategies for atlas-based image segmentation with application to confocal microscopy images of bee brains. NeuroImage 21, 1428–1442.

Ronneberger, O., Fischer, P., and Brox, T. (2015). U-net: Convolutional networks for biomedical image segmentation. In Lecture Notes in Computer Science, pages 234–241. Springer International Publishing.

Ruppert, D., Wand, M. P., and Carroll, R. J. (2003). Semiparametric Regression. Cambridge University Press.

Sandor, S. and Leahy, R. (1997). Surface-based labeling of cortical anatomy using a deformable atlas. IEEE Transactions on Medical Imaging 16, 41–54.

Sarracanie, M. and Salameh, N. (2020). Low-field MRI: How low can we go? a fresh view on an old debate. Frontiers in Physics 8,.

Satterthwaite, T. D., Connolly, J. J., Ruparel, K., Calkins, M. E., Jackson, C., Elliott, M. A., Roalf, D. R., Hopson, R., Prabhakaran, K., Behr, M., Qiu, H., Mentch, F. D., Chiavacci, R., Sleiman, P. M., Gur, R. C., Hakonarson, H., and Gur, R. E. (2016). The philadelphia neurodevelopmental cohort: A publicly available resource for the study of normal and abnormal brain development in youth. NeuroImage 124, 1115–1119.

Seydel, K. B., Kampondeni, S. D., Valim, C., Potchen, M. J., Milner, D. A., Muwalo, F. W., Birbeck, G. L., Bradley, W. G., Fox, L. L., Glover, S. J., Hammond, C. A., Heyderman, R. S., Chilingulo, C. A., Molyneux, M. E., and Taylor, T. E. (2015). Brain swelling and death in children with cerebral malaria. New England Journal of Medicine 372, 1126–1137.

Shattuck, D. W., Sandor-Leahy, S. R., Schaper, K. A., Rottenberg, D. A., and Leahy, R. M. (2001). Magnetic resonance image tissue classification using a partial volume model. NeuroImage 13, 856–876.

Sheth, K. N., Mazurek, M. H., Yuen, M. M., Cahn, B. A., Shah, J. T., Ward, A., Kim, J. A., Gilmore, E. J., Falcone, G. J., Petersen, N., Gobeske, K. T., Kaddouh, F., Hwang, D. Y., Schindler, J., Sansing, L., Matouk, C., Rothberg, J., Sze, G., Siner, J., Rosen, M. S., Spudich, S., and Kimberly, W. T. (2020). Assessment of brain injury using portable, low-field magnetic resonance imaging at the bedside of critically ill patients. JAMA Neurology

Shi, F., Wang, L., Dai, Y., Gilmore, J. H., Lin, W., and Shen, D. (2012). LABEL: Pediatric brain extraction using learning-based meta-algorithm. NeuroImage 62, 1975–1986.

Shinohara, R. T., Sweeney, E. M., Goldsmith, J., Shiee, N., Mateen, F. J., Calabresi, P. A., Jarso, S., Pham, D. L., Reich, D. S., and Crainiceanu, C. M. (2014). Statistical normalization techniques for magnetic resonance imaging. NeuroImage: Clinical 6, 9–19.

Sled, J., Zijdenbos, A., and Evans, A. (1998). A nonparametric method for automatic correction of intensity nonuniformity in MRI data. IEEE Transactions on Medical Imaging 17, 87–97.

Smith, S. M. (2002). Fast robust automated brain extraction. Human Brain Mapping 17, 143–155.

Tohka, J., Krestyannikov, E., Dinov, I., Graham, A., Shattuck, D., Ruotsalainen, U., and Toga, A. (2007). Genetic algorithms for finite mixture model based voxel classification in neuroimaging. IEEE Transactions on Medical Imaging 26, 696–711.

Tustison, N. J., Avants, B. B., Cook, P. A., Zheng, Y., Egan, A., Yushkevich, P. A., and Gee, J. C. (2010). N4itk: Improved n3 bias correction. IEEE Transactions on Medical Imaging 29, 1310–1320.

Valcarcel, A. M., Linn, K. A., Vandekar, S. N., Satterthwaite, T. D., Muschelli, J., Calabresi, P. A., Pham, D. L., Martin, M. L., and Shinohara, R. T. (2018). MIMoSA: An automated method for intermodal segmentation analysis of multiple sclerosis brain lesions. Journal of Neuroimaging 28, 389–398.

Valverde, S., Cabezas, M., Roura, E., González-Villà, S., Pareto, D., Vilanova, J. C., Ramió-Torrenta, L., Rovira, A., Oliver, A., and Lladó, X. (2017). Improving automated multiple sclerosis lesion segmentation with a cascaded 3d convolutional neural network approach. NeuroImage 155, 159–168.

Vovk, U., Pernus, F., and Likar, B. (2007). A review of methods for correction of intensity inhomogeneity in MRI. IEEE Transactions on Medical Imaging 26, 405–421.

Wang, H., Suh, J. W., Das, S. R., Pluta, J. B., Craige, C., and Yushkevich, P. A. (2013). Multi-atlas segmentation with joint label fusion. IEEE Transactions on Pattern Analysis and Machine Intelligence 35, 611–623.

World Health Organization (2020). World malaria report 2020: 20 years of global progress and challenges.

Yushkevich, P. A., Piven, J., Cody Hazlett, H., Gimpel Smith, R., Ho, S., Gee, J. C., and Gerig, G. (2006). User-guided 3D active contour segmentation of anatomical structures: Significantly improved efficiency and reliability. Neuroimage 31, 1116–1128.

Zhang, Y., Brady, M., and Smith, S. (2001). Segmentation of brain MR images through a hidden markov random field model and the expectation-maximization algorithm. IEEE Transactions on Medical Imaging 20, 45–57.

