## Supplemental Figures and Tables for "Automated Analysis of Low-Field Brain MRI in Cerebral Malaria"

by Tu *et al.*

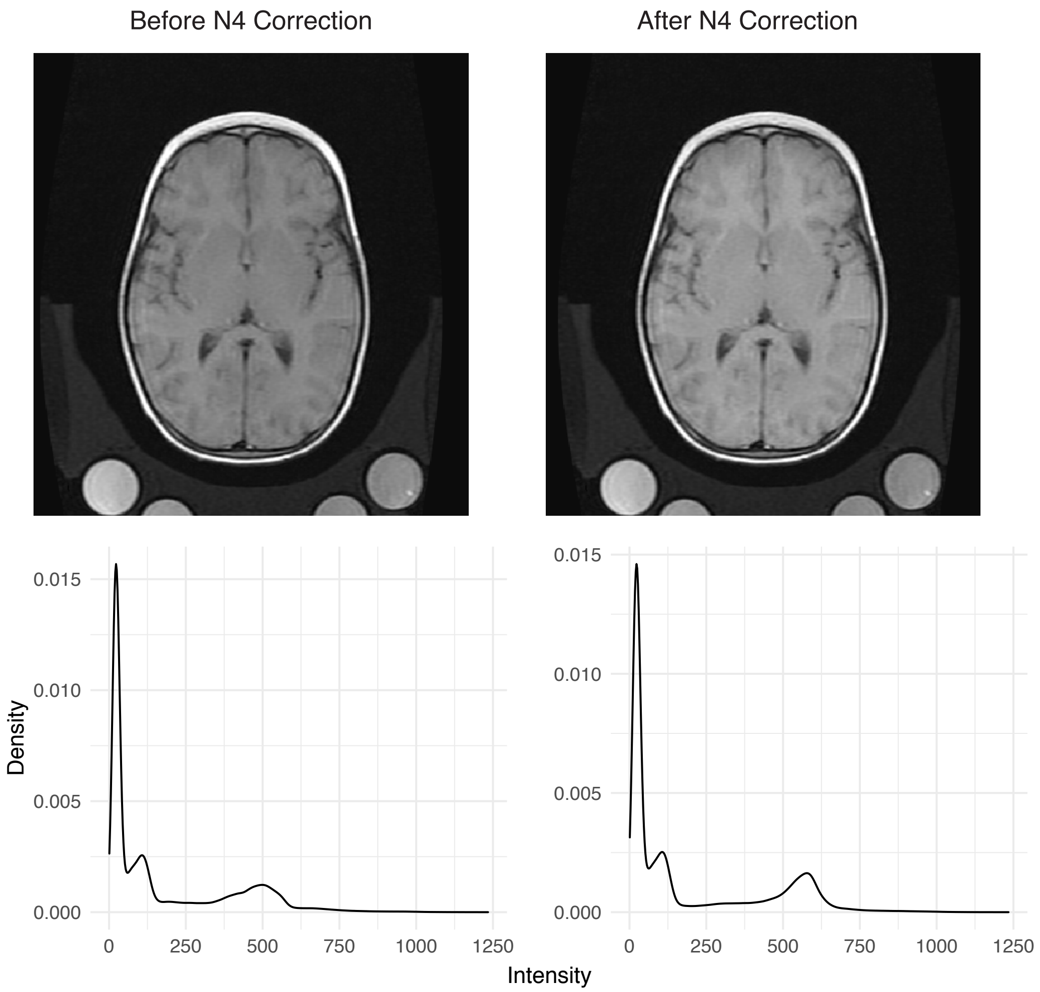


*Web Figure 1.* The effect of N4 bias correction (Tustison, Cook, & Gee, 2011) on a T1-weighted image and the corresponding intensity distributions. (Left column) The raw T1 image before N4 bias correction and the intensity histogram of non-zero voxels. (Right column) The same brain scan after N4 correction.

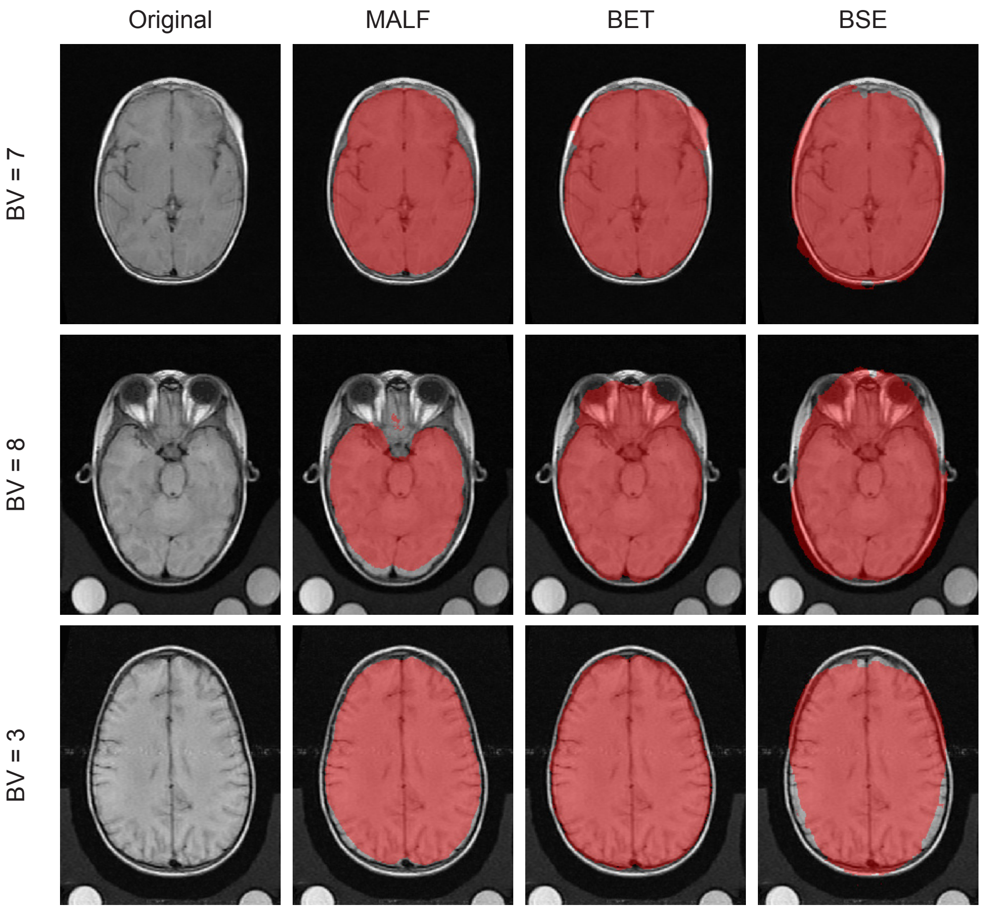


*Web Figure 2.* In 0.35 Tesla MRI images, a multi-atlas label fusion (MALF) method (Doshi, Erus, Ou, Gaonkar, & Davatzikos, 2013) performs better in brain segmentation compared to standard methods that work well on high-resolution images. In three selected participants with BV = 3, 7, and 8, an axial slice is shown with labeled brain voxels in red. We compare the performance of MALF against two standard methods, the Brain Extraction Tool (BET) (Smith, 2002) and the Brain Surface Extractor (BSE) from BrainSuite (Sandor & Leahy, 1997). Overall, we find that MALF has the more accurate and conservative segmentation, while BST and BSE perform worse in the inferior slices near the optic nerve, and in the superior slices near the top of the head.

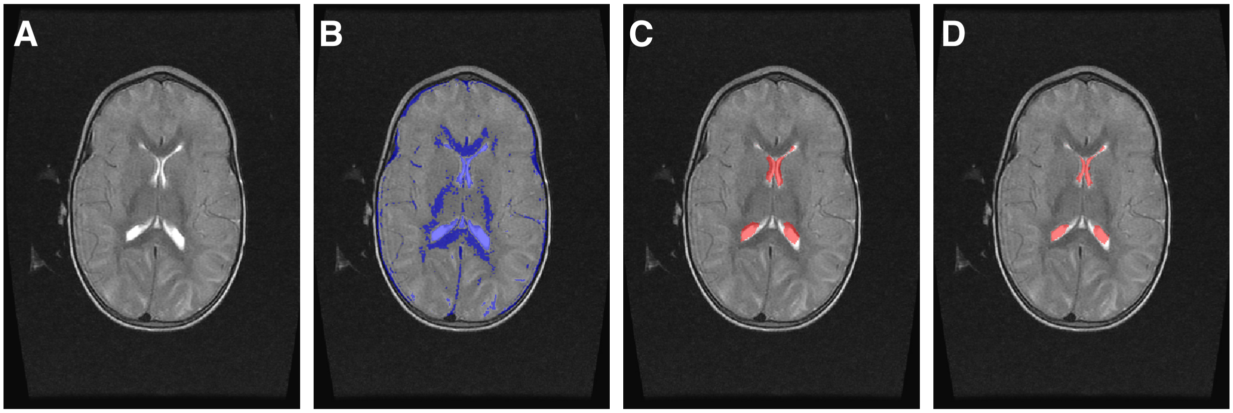


*Web Figure 3.* We identified ventricular CSF voxels by taking the intersection of the CSF mask from FSL FAST and the ventricle mask from the OASIS atlases. (A) The original T2 image. (B) The CSF segmentation from FSL FAST. (C) Ventricle segmentation obtained by multi-atlas label fusion on adult OASIS ventricle atlases. (D) The intersection of masks from Panels B and C representing the ventricular CSF mask. By taking the intersection, segmentation errors observed in the FSL FAST and OASIS segmentations are minimized.

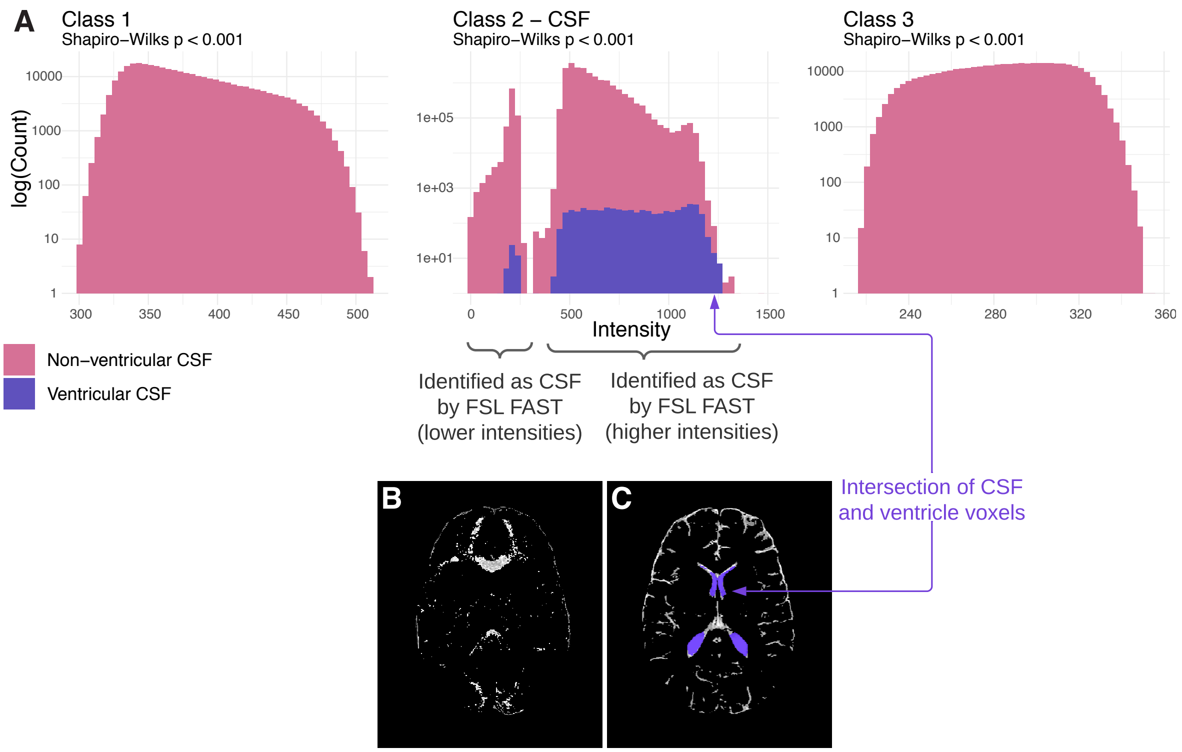


*Web Figure 4.* By using the intersection of CSF and ventricle voxels, errors from FSL FAST's tissue classification are attenuated. (A) Intensity histograms for a randomly selected participant's T2 scan, where tissue classes are determined using FSL FAST. For all tissue classes, the distribution of voxel intensities was non-Gaussian, based on the Shapiro-Wilks test performed on a sample of 6,000 voxels. In particular, the distribution of CSF (Class 2) voxel intensities was frequently bimodal. The purple bars correspond to ventricular CSF voxels, which were identified as the intersection of CSF voxels (found by FSL FAST) and ventricle voxels (found by multi-atlas label fusion of the OASIS atlases). (B) The lower component of the CSF distribution maps to voxels that are incorrectly identified as CSF. (C) The upper component of the CSF distribution maps to voxels which contain both correctly identified CSF voxels. The overlaid purple regions show the voxels ultimately identified as ventricular CSF.

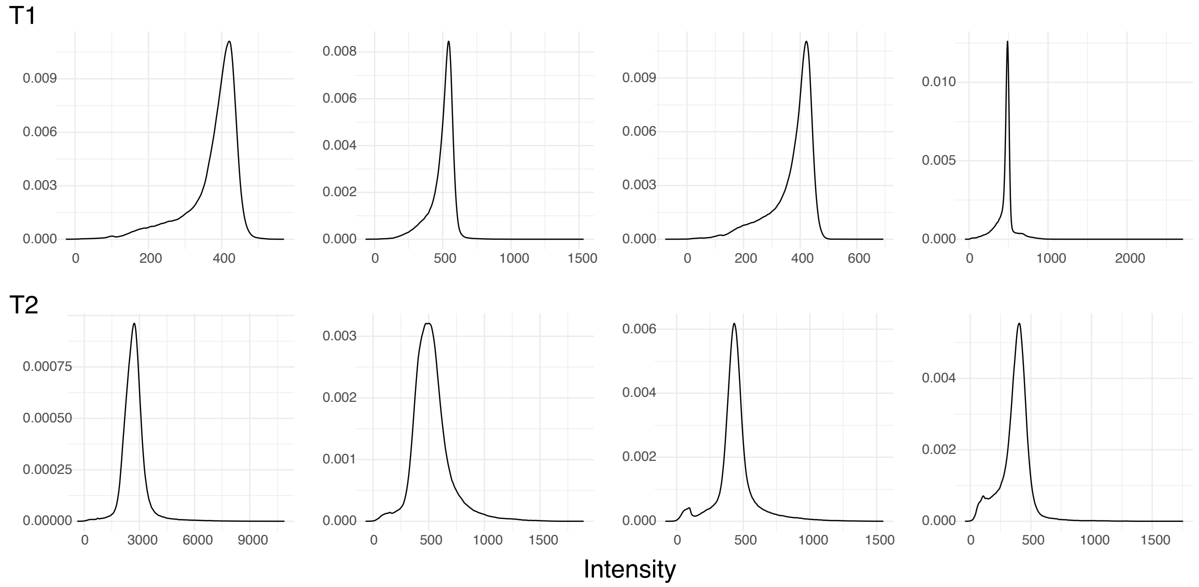


*Web Figure 5.* Densities of in-brain voxels for eight randomly selected T1 and T2 images. Between 2 and 3 peaks can be observed in each density, corresponding to the tissue types within the image. While there is no formal test to determine the true data generating process of the data, these densities support the assumption that the intensity density can be modeled as a mixture of 3 distributions.

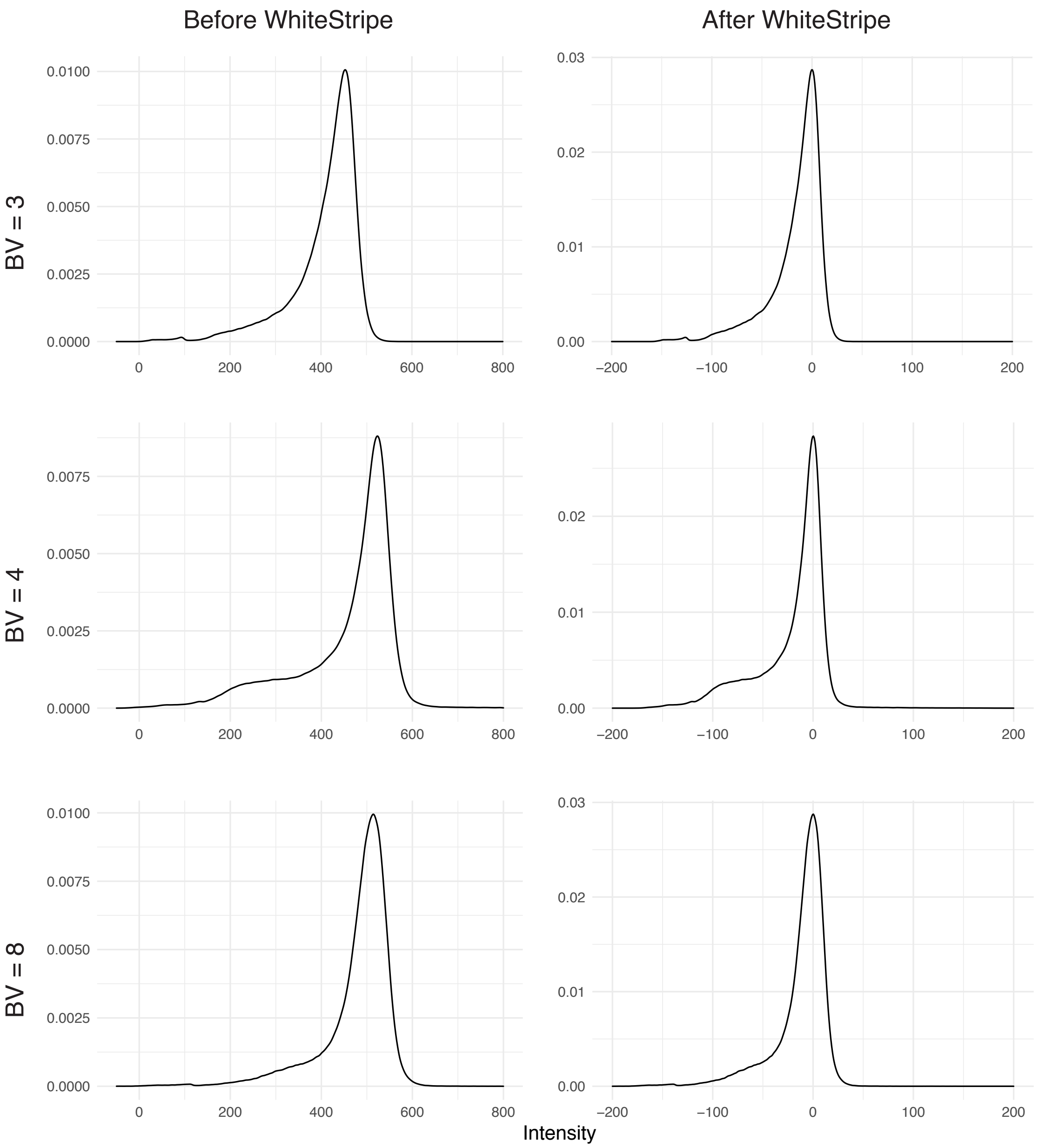


*Web Figure 6.* The effect of WhiteStripe on the intensity histogram of in-brain voxels in the T1 image for three selected participants with BV = 3, 4, and 8. After normalizing data, intensity-based features can be compared across participants. (Left column) Before WhiteStripe, the intensity distributions have different modes and spreads due to the arbitrary intensity units obtained from MRI acquisition. (Right column) After WhiteStripe, the highest peaks (corresponding to white matter in T1 images) are aligned.

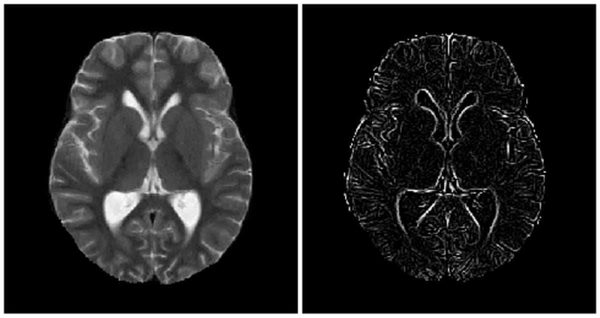

*Web Figure 7.* A T2-weighted brain MRI from a participant with median BV score of 2.5. (Left) T2 axial slice. (Right) Hessian filter highlighting ridges and gyri within the eroded brain mask.

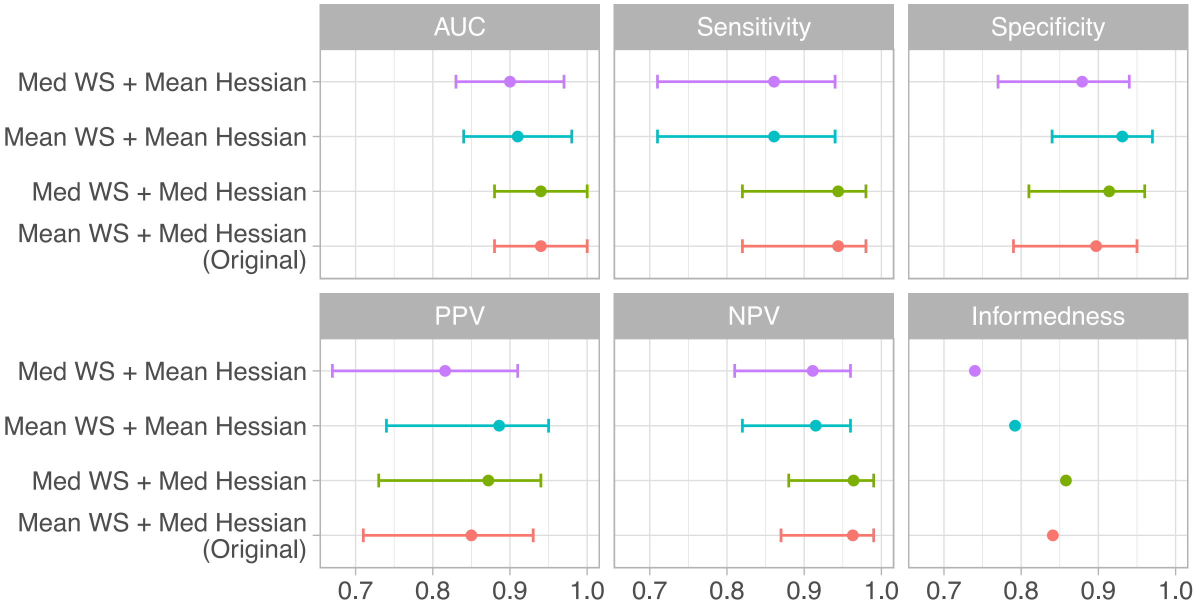


*Web Figure 8.* Performance accuracy of the prediction model is robust to different biomarker specifications. We considered either the mean or median of the WhiteStripe (WS) intensities and Hessian filter, in addition to the BPF, in predicting cases with BV $\geq$ 7. Using nested cross-validation, we find that the AUC, sensitivity, specificity, PPV, and NPV are not significantly different between the 4 combinations.

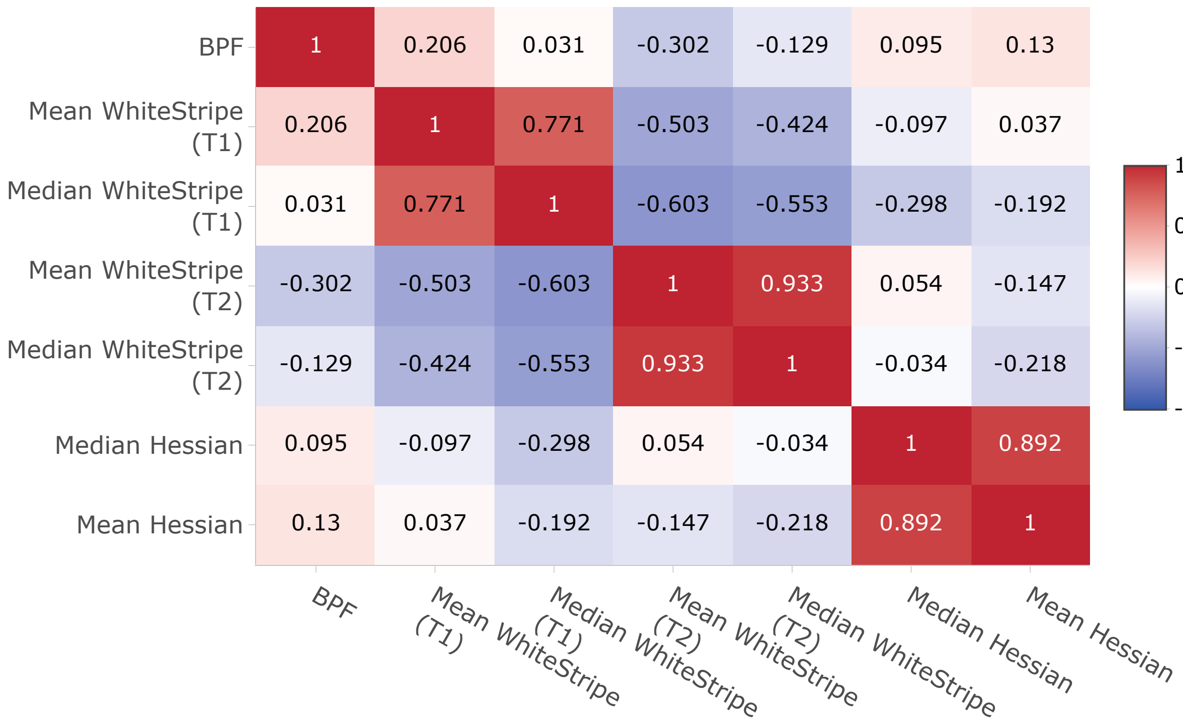


*Web Figure 9.* Correlation heatmap between biomarkers. Within each biomarker, the mean and median values were highly correlated. The WhiteStripe values from T1 and T2 were also inversely correlated, as expected.

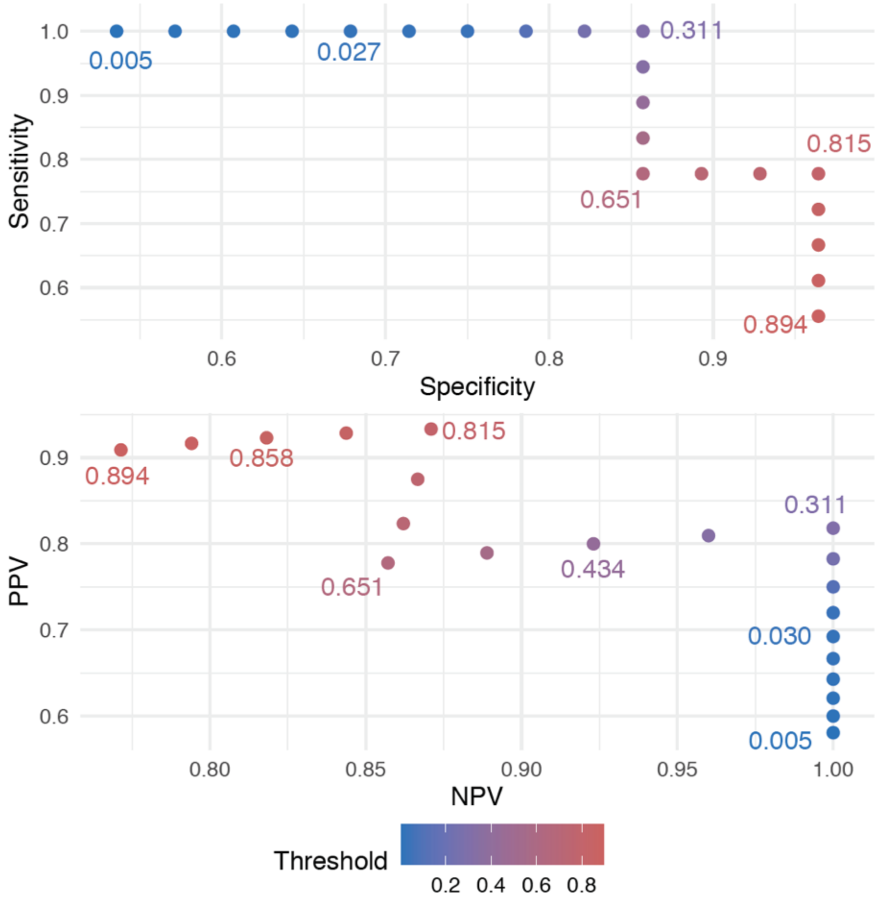


*Web Figure 10.* Given the primary model (1), the choice of threshold affects the sensitivity, specificity, positive predictive value (PPV), and negative predictive value (NPV) of the predicted outcome. We considered thresholds for which the sensitivity and specificity were higher than 50%. We found that a wide range of thresholds, ranging from around 0.3 to 0.8, yielded a favorable compromise between all four measures, while guaranteeing that each is higher than 75%.

| | Dependent variable: |
|---|---|
| | *BV Score* |
| $\gamma_{.1}$(Brain Parenchymal Fraction) | -72.059 |
| | (-170.977, 26.858) |
| $\gamma_{.2}$ (Mean WhiteStripe – T1) | 0.064** |
| | (0.015, 0.113) |
| $\gamma_{.3}$ (Mean WhiteStripe – T2) | -0.275*** |
| | (-0.329, -0.222) |
| $\gamma_{.4}$ (Sulcal Effacement – T2) | -0.094** |
| | (-0.181, -0.007) |
| Constant | 77.961 |
| | (-20.523, 176.444) |
| Observations | 94 |
| R2 | 0.677 |
| Adjusted R2 | 0.662 |
| Residual Std. Error | 1.130 (df = 89) |
| F Statistic | 46.577*** (df = 4; 89) |
| *Note:* | *p<0.1; **p<0.05; ***p<0.01 |

*Web Table 1.* Estimated coefficients from the linear model predicting BV score directly. This model was fit on the combined training and testing data (n = 94). While the main clinical outcome of interest was the identification of cases with highly increased brain volume (BV), motivating the application of a logistic model, we also considered predicting the BV score directly, and as a function of the same covariates as the logistic model.

| Model | BV Threshold | Sensitivity | Specificity | NPV | PPV | Youden's *J* |
|---|---|---|---|---|---|---|
| Linear | 7 | 0.95 (0.82,1) | 0.35 (0.1,0.59) | 0.8 (0.26,1) | 0.7 (0.59,0.82) | 0.3 (0,0.62) |
| **Logistic** | **7** | **0.89 (0.75,1)** | **0.92 (0.68,1)** | **0.85 (0.65,1)** | **0.95 (0.81,1)** | **0.82 (0.66,0.97)** |
| Linear | 6 | 0.97 (0.85,1) | 0.74 (0.48,1) | 0.98 (0.87,1) | 0.77 (0.51,1) | 0.71 (0.35,1) |
| Logistic | 6 | 0.81 (0.69,0.93) | 0.9 (0.61,1) | 0.84 (0.64,1) | 0.83 (0.36,1) | 0.71 (0.4,1) |

*Web Table 2.* Prediction performance of the thresholded linear regression model compared to the logistic regression model. All models were fit on the full set of covariates. The main model in (1) is highlighted in bold. For determining cases with BV $\geq 7$, the linear regression model performs comparably to the logistic model in sensitivity, but has lower specificity, pointing to a higher rate of false negatives.

| Threshold | Sensitivity | Specificity | PPV | NPV |
|---|---|---|---|---|
| 0.005 | 1 | 0.54 | 0.58 | 1 |
| 0.007 | 1 | 0.57 | 0.6 | 1 |
| 0.009 | 1 | 0.61 | 0.62 | 1 |
| 0.018 | 1 | 0.64 | 0.64 | 1 |
| 0.027 | 1 | 0.68 | 0.67 | 1 |
| 0.03 | 1 | 0.71 | 0.69 | 1 |
| 0.036 | 1 | 0.75 | 0.72 | 1 |
| 0.118 | 1 | 0.79 | 0.75 | 1 |
| 0.251 | 1 | 0.82 | 0.78 | 1 |
| 0.311 | 1 | 0.86 | 0.82 | 1 |
| 0.333 | 0.94 | 0.86 | 0.81 | 0.96 |
| 0.434 | 0.89 | 0.86 | 0.8 | 0.92 |
| 0.561 | 0.83 | 0.86 | 0.79 | 0.89 |
| 0.651 | 0.78 | 0.86 | 0.78 | 0.86 |
| 0.73 | 0.78 | 0.89 | 0.82 | 0.86 |
| 0.773 | 0.78 | 0.93 | 0.88 | 0.87 |
| 0.815 | 0.78 | 0.96 | 0.93 | 0.87 |
| 0.844 | 0.72 | 0.96 | 0.93 | 0.84 |
| 0.858 | 0.67 | 0.96 | 0.92 | 0.82 |
| 0.88 | 0.61 | 0.96 | 0.92 | 0.79 |
| 0.894 | 0.56 | 0.96 | 0.91 | 0.77 |

*Web Table 3.* Given the primary model (1), the choice of threshold affects the sensitivity, specificity, positive predictive value (PPV), and negative predictive value (NPV) of the predicted outcome. We considered thresholds for which the sensitivity and specificity were higher than 50%. We found that a wide range of thresholds, ranging from around 0.3 to 0.8, yielded a favorable compromise between all four measures, while guaranteeing that each is higher than 75%.

| Outcome | Covariates | Threshold on Predicted Probability | Sensitivity | Specificity |
|---|---|---|---|---|
| **BV ≥ 7** | **All Biomarkers** | **0.711** | **0.944** | **0.929** |
| BV ≥ 7 | Sulcal Only | 0.731 | 0.944 | 0.929 |
| BV ≥ 6.5 | All Biomarkers | 0.711 | 0.895 | 0.926 |
| BV ≥ 6.5 | Sulcal Only | 0.731 | 0.895 | 0.926 |
| BV ≥ 6 | All Biomarkers | 0.806 | 0.957 | 0.913 |
| BV ≥ 6 | Sulcal Only | 0.598 | 0.870 | 0.957 |

*Web Table 4.* We can identify specific thresholds of the predicted probably to optimize sensitivity and specificity. For each model, we considered a set of thresholds, where predicted probabilities greater than the threshold are interpreted as severe cases, and non-severe otherwise. Then, the final threshold was chosen such that the sensitivity and specificity were maximized, in that order, and with the restriction that specificity be greater than 90%. The primary model in (1) is shown in bold.

| Outcome | Covariates | Train AUC | Train AUC (95% CI) | Test AUC | Test AUC (95% CI) |
|---|---|---|---|---|---|
| **BV ≥ 7** | **Full Set** | **0.95** | **[0.90,1.00]** | **0.88** | **[0.80,0.96]** |
| BV ≥ 7 | Sulcal Only | 0.96 | [0.91,1.00] | 0.88 | [0.80,0.96] |
| BV ≥ 6.5 | Full Set | 0.92 | [0.86,0.98] | 0.81 | [0.72,0.91] |
| BV ≥ 6.5 | Sulcal Only | 0.94 | [0.88,1.00] | 0.82 | [0.73,0.91] |
| BV ≥ 6 | Full Set | 0.93 | [0.88,0.98] | 0.86 | [0.78,0.94] |
| BV ≥ 6 | Sulcal Only | 0.76 | [0.66,0.86] | 0.84 | [0.75,0.92] |

*Web Table 5.* Our model achieves good performance on external validation data. Each row corresponds to a model that was trained on the complete original data ($n$ = 94), with AUC calculated using *unnested* 5-fold cross-validation (since the inner cross-validation loop was used to determine the optimal cutoff). Each model was tested on the new set of validation data obtained from an ongoing study of brain swelling in CM ($n$ = 90). Overall, the model performed well on the new data, albeit with slightly decreased accuracy compared to the original data in 5 out of 6 models.
